# Cardiac competence of the paraxial head mesoderm fades concomitant with a shift towards the head skeletal muscle programme

**DOI:** 10.1101/759506

**Authors:** Afnan Alzamrooni, Petra Mendes Vieira, Nicoletta Murciano, Matthew Wolton, Frank R. Schubert, Samuel C. Robson, Susanne Dietrich

## Abstract

The vertebrate head mesoderm provides the heart, the great vessels, some smooth and most head skeletal muscle, in addition to parts of the skull. It has been speculated that the ability to generate cardiac and smooth muscle is the evolutionary ground-state of the tissue. However, whether indeed the entire head mesoderm has generic cardiac competence, how long this may last, and what happens as cardiac competence fades, is not clear.

Bone morphogenetic proteins (Bmps) are known to promote cardiogenesis. Using 41 different marker genes in the chicken embryo, we show that the paraxial head mesoderm that normally does not engage in cardiogenesis has the ability to respond to Bmp for a long time. However, Bmp signals are interpreted differently at different time points. Up to early head fold stages, the paraxial head mesoderm is able to read Bmps as signal to engage in the cardiac programme; the ability to upregulate smooth muscle markers is retained slightly longer. Notably, as cardiac competence fades, Bmp promotes the head skeletal muscle programme instead. The switch from cardiac to skeletal muscle competence is Wnt-independent as Wnt caudalises the head mesoderm and also suppresses Msc-inducing Bmp provided by the prechordal plate, thus suppressing both the cardiac and the head skeletal muscle programmes.

Our study for the first time suggests a specific transition state in the embryo when cardiac competence is replaced by skeletal muscle competence. It sets the stage to unravel the cardiac-skeletal muscle antagonism that is known to partially collapse in heart failure.

**Summary statement:** The head mesoderm has generic cardiac competence until early head fold stages. Thereafter, cardiac competence fades in the paraxial region, and Bmp promotes head skeletal muscle programmes instead of cardiac programmes.

## 1. Introduction

The vertebrate head mesoderm is the first embryonic mesoderm type to emerge during gastrulation (Camp et al., 2012; Garcia-Martinez and Schoenwolf, 1993). It delivers the heart (reviewed in (Meilhac and Buckingham, 2018; Sendra et al., 2021; Spater et al., 2014)), the great vessels (Paffett-Lugassy et al., 2013; Wang et al., 2017), the smooth muscle collar of the cardiac outflow (Wang et al., 2017), genuine craniofacial and oesophageal skeletal muscles (reviewed in (Schubert et al., 2018)) and parts of the skull ((Jandzik et al., 2015); reviewed in (Kuratani, 2005)), with cardiac cells originating from the lateral, and skeletal muscle and also bone/cartilage mainly from the paraxial region of the tissue (reviewed in (Evans and Noden, 2006)). The primitive heart, blood and vascular system are the first fully functional organs in the embryo, and defects in their development are incompatible with life. In recent years, the head mesoderm has received quite some attention since a subpopulation of cardiogenic cells, often referred to as second or secondary heart field (SHF) cells and located in the pharyngeal arches at the junction to the primitive heart, seem to be multipotent cardiac-skeletal muscle-vascular endothelial precursor or stem cells (reviewed in (Sendra et al., 2021)). When these cells are recruited into the primitive heart, they deliver the mature in-and outflow tract, in mammals and birds also encompassing the right ventricle. This process is essential as compromised SHF formation causes cardiac birth defects (reviewed in (Houyel and Meilhac, 2021; Morton et al., 2022)). Notably, SHF cells integrate perfectly into the developing heart and beat according to the rhythm set by the pre-existing cells. This is in stark contrast to cells transplanted into an adult heart to replace cells lost after a heart attack: here, cells struggle to integrate, and their autonomous beating may cause potentially fatal arrhythmias (Cui et al., 2018; Scuderi and Butcher, 2017; Spater et al., 2014). In a similar vein, the embryo is able to allocate head mesodermal cells to various conflicting developmental programmes. Yet in a failing heart, cells partially lose their cardiomyocyte identity and erroneously express skeletal muscle genes (reviewed in (Oh et al., 2019)). Thus, sustained efforts aim to understand the properties and developmental programmes of cells in the embryonic head mesoderm in order to apply this understanding to diagnosis and treatment of disease.

Transcriptome analyses and grafting experiments showed that the head mesoderm is distinct from the mesoderm in the trunk that does not normally deliver cardiomyocytes and that forms skeletal muscle using distinct programmes (reviewed in (Schubert et al., 2018)). Comparative analyses between vertebrates and invertebrate chordates indicated that this head-trunk distinction is basic to chordates. Notably, chordates originally lacked mesoderm-derived cartilage and bones, and skeletal (voluntary) muscle served the purpose of locomotion and formed from the trunk paraxial mesoderm (somites). Contractile tissue in the head existed in form of pharyngeal smooth muscle and cardiac muscle placed upstream of the vessels supplying the pharyngeal basket/pharyngeal arches (reviewed in (Poelmann and Gittenberger-de Groot, 2019; Simões-Costa et al., 2005). Recent analyses of the core regulatory complexes controlling muscle formation suggested that the striated organisation of cardiac muscle originated from a recruitment of downstream skeletal muscle effector genes into pre-existing smooth muscle programme (Brunet et al., 2016). However, with the acquisition of an active, predatory lifestyle, vertebrates seem to have recruited further aspects of the skeletal muscle programme into the head mesoderm, including the MyoD-type core regulators of the voluntary/skeletal muscle programme, establishing a voluntary control over the cranial openings, food uptake and in humans, speech. Given this sequence of events, we wondered whether the vertebrate paraxial head mesoderm (PHM) which does not normally partake in cardiogenesis might initially be bestowed with cardiac competence and might at some stage in development switch from cardiac to skeletal muscle competence. The aim of this study was to investigate this.

Cells fated to contribute to the heart are specified as cardiogenic cells, but not yet determined, when they leave the primitive streak ((Lopez-Sanchez et al., 2009; Wang et al., 2013); reviewed in (Sendra et al., 2021)). Migration to their target area lateral to the developing neural plate is controlled by bone morphogenetic proteins (Bmp; ((Song et al., 2014)). Moreover, Bmps are essential for cardiogenic cells to realise their potential and differentiate into cardiomyocytes (reviewed in (Cui et al., 2018; Spater et al., 2014; van Wijk et al., 2007)). Bmps drive cardiogenesis indirectly by suppressing the proliferation-promoting function of fibroblast growth factors (Hutson et al., 2010; Tirosh-Finkel et al., 2010). They also control cardiogenesis directly, with Smad1/5/8- Smad4 complexes transactivating genes encoding key cardiac transcription factors such as Isl1, Nkx2.5, Gata4 and Tbx2 (Brown et al., 2004; Hami et al., 2011; Liberatore et al., 2002; Lien et al., 2002; Shirai et al., 2009; Si et al., 2014). Before and during the formation of the primitive heart, the Bmp ohnologues Bmp2 and Bmp4 are expressed in the lateral and extraembryonic mesoderm, its overlying ectoderm and underlying pharyngeal endoderm and the neural folds. Bmp2 continues to be expressed in the cardiac inflow tract, Bmp4 in the SHF adjacent to the heart. The heart expresses Bmp7 throughout, the ventricle expresses Bmp10. Thus, the lateral head mesoderm is under the influence of Bmp all the time. In contrast, the PHM is exposed to the Bmp inhibitors Noggin and Chordin delivered by Hensen’s node and the notochord in the midline of the embryo ((Bothe et al., 2011), http://geisha.arizona.edu/geisha/; for Bmp phylogeny see (Hinck et al., 2016; Huminiecki et al., 2009)). Moreover, various studies showed that exposure to Bmp can trigger the expression of cardiac markers in the PHM (Andree et al., 1998; Bothe et al., 2011; Schlange et al., 2000; Schultheiss et al., 1997; Tirosh-Finkel et al., 2006; von Scheven et al., 2006a; Yamada et al., 2000). Yet, Bmp7 is expressed in the early prechordal plate and later, in the pharyngeal arches (von Scheven et al., 2006a). It thus is in the position to signal to the developing branchiomeric skeletal muscles. Moreover, Bmp signalling has been shown to activate the expression of *Msc* (Bothe et al., 2011), a gene in the head involved in the activation of the first MyoD family member to be expressed, namely *Myf5* and *MyoD* ((Moncaut et al., 2012), reviewed in (Schubert et al., 2018)). We therefore reasoned that a strict exposure regime to Bmp would reveal the initial and any shift in the cellular competence of the PHM. Our study shows that this indeed was the case: differing, stage-dependent responses to Bmp indicate a shift from cardiac to skeletal muscle competence in the PHM.

β-Catenin-mediated Wnt signalling plays multiple roles during cardiac development (reviewed in (Gessert and Kühl, 2010). Wnt signalling is required for gastrulation, and hence without Wnt, cardiogenic cells cannot form (Kraus et al., 2016). Wnt signals deflect the early gastrulating cells from the primitive streak and hence contribute to the congregation of cardiogenic cells in the lateral head mesoderm (Song et al., 2014; Yue et al., 2008). However, prolonged Wnt signals also contribute to the caudalisation of the gastrulating mesoderm, thus establishing the trunk programmes and suppressing cardiogenesis (Marvin et al., 2001). To escape this inhibition, cardiac precursors are bestowed with the expression of Wnt inhibitors such as Szl (also called Sfrp2- or Sfrp5 related; (Wittler et al., 2008), reviewed in (Wu et al., 2020)). Previous studies proposed that an increase in paraxial β-Catenin-dependent Wnt signalling may suppress the ability of the PHM to engage in cardiogenesis (Tzahor and Lassar, 2001). We therefore also tested whether Wnt signalling may promote the switch from cardiac to skeletal muscle competence in the PHM. However, this was not the case: Wnt suppressed the onset of the head skeletal muscle programme, both by caudalising the PHM and by suppressing *Msc*-inducing Bmp production from the prechordal plate, suggesting that the switch in developmental competence is intrinsic to the PHM.

## 2. Material and Methods

### 2.1. Chicken embryo culture, staging and harvesting

Wildtype fertilised chicken eggs were obtained from Medeggs Ltd. (Norfolk, UK), eggs from GFP- and TdTomato-expressing chicken were obtained from the National Avian Research Facility (https://www.ed.ac.uk/roslin/national-avian-research-facility). Eggs were incubated at 38.5°C in a humidified incubator (LMS) and staged according to (Hamburger and Hamilton, 1951). For bead and GFP-tissue implantation experiments at HH5-10, embryos were cultured as ‘EC-cultures’ on filter rings as described by Chapman and Bothe (Bothe et al., 2011; Chapman et al., 2001); experiments at HH13/14 were performed in ovo. For the RNAseq screen, HH7/8 embryos were set up as Cornish Pasty cultures as described in (Dupé and Lumsden, 2001). Embryos that received beads or tissue grafts were harvested in 4% PFA.

### 2.2. Recombinant proteins and bead preparation

Recombinant Bmp proteins from R&D Systems/Biotechne (Bmp2: 355-BM-10, Bmp4: 314-BP-10, Bmp7: 354-BP-10, Bmp10: 2926-BP-025) were reconstituted in 4mM HCl supplemented with 0.1% bovine serum albumin at a concentration of 1mg/ml or 0.5mg/ml. Affi-Gel blue agarose beads (BioRad) and heparin-coated acrylic beads (Sigma) suspended in sterile water were added to obtain 0.19μM, 1.28μM, 6.41μM or 19.23μM dilutions. Wnt3a protein from R&D Systems/Biotechne (5036- WN-010) was reconstituted in PBS with 0.1% bovine serum albumin at 0.4mg/ml; beads suspended in sterile water were added to obtain 5.35-10.7μM dilutions. No difference in phenotypes for the different Wnt3a concentrations were observed with the most sensitive marker *Cyp26C1*, and the experiments were then carried out with 8.56μM Wnt3a and the data combined. The beads were soaked in these solutions for an hour on ice, then washed in saline before grafting. Control beads were soaked in 0.1% BSA in saline.

### 2.3. Bead implantation

Beads were grafted into the paraxial head mesoderm (PHM) of HH5/6, HH7/8 and HH9/10 embryos on EC culture plates and HH13/14 embryos in ovo, using flame-sharpened tungsten needles as described in (Bothe et al., 2011; von Scheven et al., 2006a). For HH5/6 and HH7/8, embryos were accessed from the ventral side; HH9/10 embryos were accessed from the dorsal side. Unless stated otherwise, beads were implanted into the central PHM as shown in Supplementary Figure SF1. Beads for the lateral/cardiogenic head mesoderm were implanted into EC-culture embryos from the ventral side. The embryos were incubated for 6 hours, during which time they progressed to the next stage of development. For the time course experiment, embryos were incubated for 2, 4 and 6 hours. To allow for the response of indirect targets further downstream in regulatory cascades and for cardiac differentiation, embryos were also incubated overnight. All n-numbers are collated in Supplementary Table ST1.

### 2.4. GFP-tissue implantation

GFP-or TdTomato-expressing donor embryos were mounted on filter rings, washed in saline and pinned down on Sylgard-bottomed Petri dishes as described in (Alvares et al., 2008). To extract the HH5/6 and 7/8 PHM and HH5/6 cardiac mesoderm, donors were ventral side up. Using a glass capillary mounted on an aspirator, a small drop of 1mg/ml Dispase (Sigma) in DMEM was delivered to the to-be-excised region, then the endoderm was peeled off using flame-sharpened, angled 0.1mm tungsten needles. Using these needles, the PHM was cut free from the region inside the neural plate territory, the lateral/cardiac mesoderm was excised from the territory lateral to the neural plate. The tissue was transferred into saline with a further glass capillary and aspirator to wash off excess Dispase, cut into smaller fragments, and transferred to the hosts set up as EC-cultures. In the host, a small incision was made at the desired location. This released tissue tension, delivering an opening to fit in the graft.

To recover the HH9/10 PHM, the donor was pinned down dorsal side up, and Dispase was applied. Using tungsten needles, an incision was made into the surface ectoderm parallel to the neural tube, and the ectoderm plus adhering neural crest cells were peeled away. The PHM was eased away from the neural tube medially and from the surface ectoderm and lateral mesoderm laterally, cut free rostrally and, with a distance to the 1^st^ somite, caudally, and lifted out of the slit. It was washed, cut and transferred as described before.

To recover the HH13/14 PHM, the embryo was pinned down, with pins holding the head in place. Dispase was delivered to the region dorsal to the pharyngeal arches, and the surface ectoderm plus adhering neural crest cells was removed. Using the tungsten needles, a piece of mesenchyme was retrieved, washed, and implanted as before.

To excise the HH13/14 segmental plate, the donor was pinned down ventral side up, the endoderm was removed with the help of Dispase, the segmental plate was freed from the neural tube and lateral mesoderm, cut at the dorsal and ventral end, and again, washed, cut into smaller fragments and implanted.

N-numbers for the tissue grafting experiments are collated in Supplementary Table ST2.

### 2.5. Generation of templates and probes for in situ hybridisation

Probes for in situ hybridisation and wildtype gene expression patterns are summarised in Supplementary Table ST1. Templates for probe synthesis were generated using restriction endonucleases or by PCR with plasmid-based primers; antisense probes were made using the appropriate T3/T7/SP6 RNA polymerases and the DIG labelling mix (Roche/Sigma), following the manufacturer’s instructions.

### 2.6. In situ hybridisation

In situ hybridisation was performed following the procedure of (Bothe et al., 2011). Embryos were permeabilised first in methanol overnight, then with a detergent mix (3 x 6 minutes for young embryos, 3 x 10 minutes for embryos at HH13/14). They were re-fixed in 4% PFA for 20 minutes and equilibrated in pre-hybridisation mix for 1 hour. Hybridisation was performed overnight at 65-68°C; probes were detected using the alkaline phosphatase-coupled anti-DIG antibody (1:2000, Roche/Sigma) and NBT/BCIP (Roche/Sigma). The hybridisation-and antibody-step were performed using a CEM InsituPro robot. After staining, embryos were fixed in 4% PFA overnight and stored in 80% glycerol.

### 2.7. Immunofluorescence

Immunofluorescence followed the procedure described in (Berti et al., 2015; Meireles Nogueira et al., 2015), using the mouse MF20 (1:500, DSHB), the rabbit anti-GFP (1:1000, Life technology) and the rabbit RFP (1:1000, VWR) primary antibodies, and the anti-mouse IgG+IgM (H+L) – Alexa fluor 594 or 488 and the anti-rabbit (H+L) – Alexa fluor 488 or 594 secondary antibodies (1:300, Jackson Immuno).

### 2.8. Bmp-inhibitor treatment and RNAseq screen

#### 2.8.1. Treatment of embryos in “Cornish pasty” roller-tube cultures

30 embryos at HH7-8 each were prepared for “Cornish pasty” roller-tube culture as described in (Dupé and Lumsden, 2001). Batches of 5 embryos were transferred into sterile 10ml glass vials containing 2ml of culture medium (L15 Leibovitz, 10% chicken serum, 50μg/ml gentamicin). In 3 batches (i.e. 15 embryos), the medium had been supplemented with DMSO (1:1000; negative solvent control). In the other 3 batches, 1μM LDN193189 (ALK2/3 inhibitor, MedChemExpress) had been added to the medium. The embryos were cultured in a rotary hybridisation oven at 38°C for 6 hours. Thereafter, embryos that had developed normally were collected, and the whole head region up to the 1^st^ somite was harvested in RNALater. This yielded 10-15 heads per treatment or control. The procedure was repeated three times to obtain independent biological replicates.

#### 2.8.2. RNAseq screen

From the heads collected in RNALater, total RNA was prepared using the Absolutely RNA Microprep kit (Agilent). RNA yield and quality were checked using a Bioanalyzer (Agilent). Three independent biological replicates each for the control (BioSample accession number SAMN34998568) and LDN193189 treatment (BioSample accession number SAMN34998617) were sequenced on an Illumina NextSeq500 at a depth of at least 30 million paired-end reads per sample (DeepSeq, Nottingham). After quality control to assess base calling error rates and potential adapter contamination using fastQC (Andrews, 2010), samples were trimmed to remove adapter sequences and poor quality reads using Trim Galore V0.6.4_dev (Krueger, 2012) using parameters ‘--illumina -q 20 --stringency 5 -e 0.1 --length 40 --trim-n’. Trimmed reads were mapped to the current chicken reference genome (bGalGal1.mat.broiler.GRCg7b) from Ensembl using STAR Universal RNAseq aligner V2.7.3a (Dobin et al., 2013) with parameters ‘--outSAMmultNmax 300 --outSAMstrandField intronMotif’. Read counts over unique genes were identified based on gene models from Ensembl V107, and were quantified in R (R Core Team, 2021) using the summarizeOverlaps() function in the GenomicAlignments package (Lawrence et al., 2013) using parameters ‘mode = “Union”, singleEnd ⍰=⍰FALSE, ignore.strand⍰=⍰FALSE, fragments ⍰=⍰FALSE, preprocess.reads⍰=⍰invertStrand’. p values were adjusted for multiple comparisons using the (Benjamini and Hochberg, 1995) False Discovery Rate correction.

Since we analysed whole heads with a mixed cell population, expression levels of genes specifically expressed in one cell type may be low. Likewise, expression differences between the LDN193189- treated embryos and the controls are expected to be small in the short 6-hour incubation. To ensure that the majority of experimentally verified BMP-responsive genes were included, we applied thresholds of >1.3 fold change (up or down compared to DMSO-treated), with an adjusted p value <0.2. To minimise the impact of low abundance transcripts, genes with a Fragment Per Kilobase Mapped (FPKM) value <0.1 in both conditions were removed. Of the resulting 98 BMP-dependent (i.e. downregulated after LDN19318 treatment), protein-coding genes, 4 were not automatically assigned a gene name. 3 of these could be named after synteny mapping against their paralogues. The genes were analysed for gene ontology over-representation against the human dataset using the clusterProfiler 4.0 package (Wu et al., 2021). The enrichGO() function with parameters ‘OrgDb = org.Hs.eg.db, ont = "BP", keyType = "SYMBOL", pvalueCutoff = 0.01, qvalueCutoff = 0.05, pAdjustMethod = "fdr", minGSSize = 100, maxGSSize = 100000’ was used to identify the most enriched ‘Biological Process’ GO terms.

### 2.8. Photomicroscopy

Images were captured on a Zeiss Axioskop with DIC and fluorescence optics, using a Zeiss AxioCam digital camera with ZEN light software. Confocal images were captured on a Zeiss LSM 710 microscope. Images were processed using Adobe Photoshop 6.0. To photograph the heads of embryos treated at HH13/14, embryos were bisected midsagitally, using flame-sharpened tungsten needles. The colour for MF20/green-stained TdTomato/red grafts was reversed to ease comparison with the MF20/red-stained grafts from GFP/green embryos.

## 3. Results

### 3.1. Preparatory experiments

Aim of the study was to explore the developmental competence of the paraxial head mesoderm (PHM) as revealed by its stage-dependent responses to Bmp signalling. We focused on the chicken embryo because of its large size, clear anatomy and ease of access, and because it is an established model for vertebrate, including human, head mesoderm and heart development (reviewed in (Wittig and Münsterberg, 2020)). As we previously found dynamic changes in gene expression pattern and mesodermal behaviour in the PHM between developmental stages HH5/6 and HH9/10 (Bothe et al., 2011; Bothe and Dietrich, 2006), this study), and because it has been proposed that Wnt levels triggering the β-Catenin pathway may rise in this time period to suppress cardiogenesis in the PHM (Tzahor and Lassar, 2001), we also tested whether Wnt signalling was responsible for a change in the developmental competence of the PHM. As a first step in our study we carried out a series of preparatory experiments to establish a robust and also systematic and strict experimental approach, and we established that the PHM is Bmp and Wnt sensitive during the period of investigation, i.e from early neurula stages at HH5/6 to early organogenesis stages at HH13/14. This was because earlier studies had used different ligands, different delivery methods, different treatment durations and different marker genes, thus leading to conflicting results ((Andree et al., 1998; Bothe et al., 2011; Marvin et al., 2001; Schlange et al., 2000; Schultheiss et al., 1997; Song et al., 2014; Tirosh-Finkel et al., 2006; Tzahor and Lassar, 2001; von Scheven et al., 2006a; Yamada et al., 2000; Yue et al., 2008). The similarities and differences between previous studies and our study are summarised in Supplementary Figure SF1.

#### 3.1.1. Preparatory experiments to establish a strict experimental approach

When signalling molecules are delivered by biological carriers such as expression constructs or engineered cells, there is little control over the concentrations of the ligands. We therefore used recombinant proteins loaded onto heparin-coated acrylic “white” beads or Affi-Gel blue agarose beads at defined concentrations, and we tested in Bmp-sensitive embryos at HH5/6 which concentration of ligand on which bead type may reliably trigger the upregulation of Bmp-responsive genes. We used *Isl1*, a gene expressed in the cardiac mesoderm and underlying endoderm as marker, and a 6-hours treatment regime which had been shown to be sufficient for *Isl1* induction at HH5/6 before (Bothe et al., 2011). Key n-numbers for this and the subsequent bead-grafting experiments are mentioned in the text, all n-numbers for the bead experiments are provided in Supplementary Table ST1).

Our experiments revealed that BSA-loaded control beads would not upregulate *Isl1* in the PHM (n=4, Fig.1A, blue arrowheads), and Bmp2 at 0.19 µM very slightly upregulated the gene once (n=1:7, Fig.1B; Supplementary Table ST1 and data not shown). 1.28 µM Bmp2 mildly (n=3:12), and 6.41 µM Bmp2 robustly (n= 12:16) upregulated the gene (Fig.1C,D; green arrowheads). Corresponding results were obtained for the Bmp-responsive Bmp inhibitor *Noggin* (see Supplementary Table ST1). The effect was independent of the bead type, and we continued the study with “white” beads that are better retained in the tissue.

**Figure 1.**
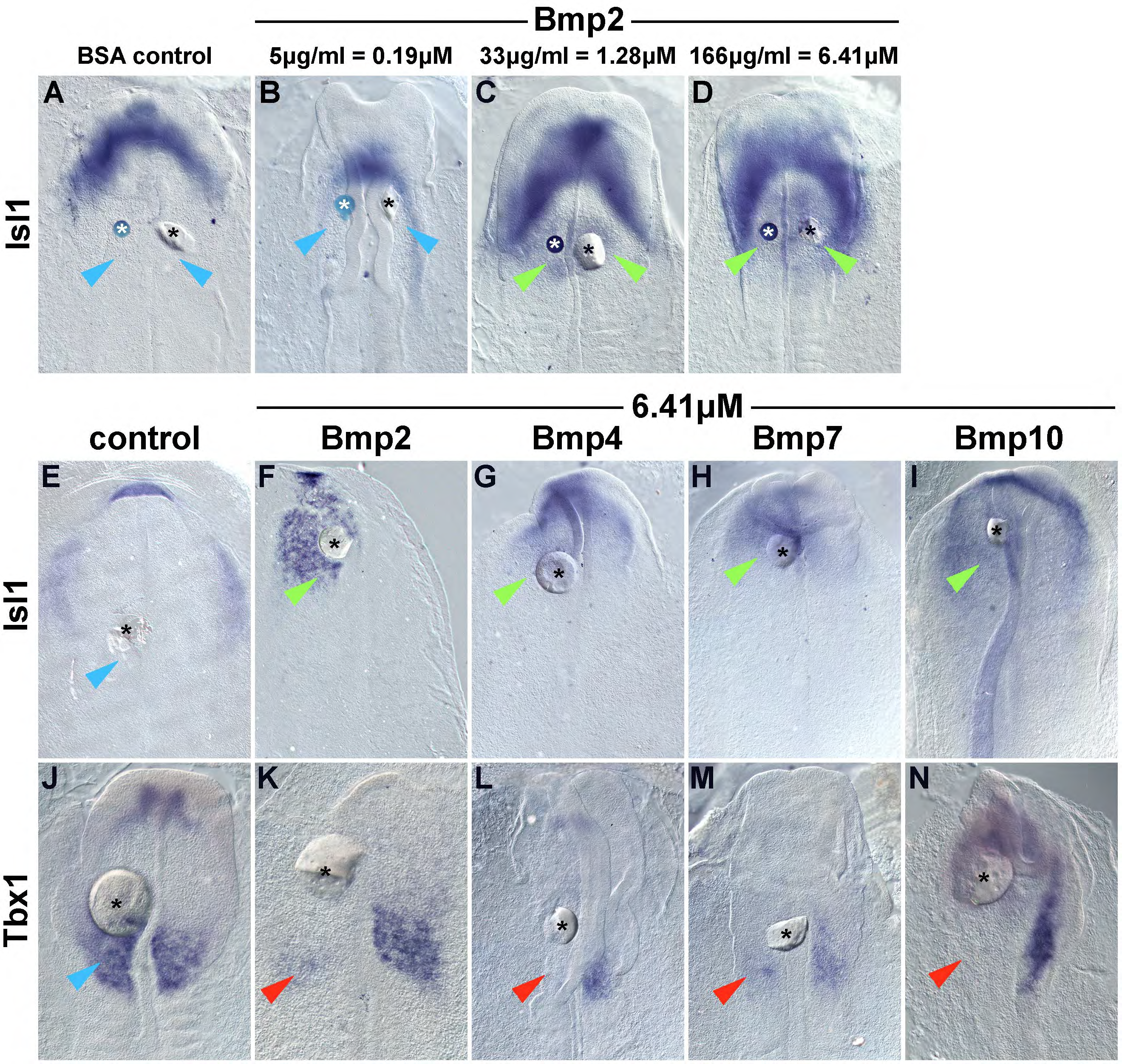
Effect of different bead types, Bmp concentrations and Bmp ligands on the HH5/6 PHM. (A-D) Dorsal views, rostral to the top, of chicken embryos that had Affi-Gel blue agarose beads (left side) or heparin-coated acrylic “white” beads (right side) implanted into their paraxial head mesoderm (PHM) at stage HH5/6. The beads are marked by asterisks. They had been loaded with BSA (control) or three different concentrations of Bmp2 as indicated on the top of the panel. After six hours of re-incubation, the embryos were analysed for the expression of *Isl1*, a marker for the cardiovascular lateral head mesoderm and the pharyngeal endoderm. Note the mild upregulation of *Isl1* with 1.28µM Bmp2 (C, green arrowheads), and the robust upregulation with 6.41µM Bmp2 (D, green arrowheads); the two bead types produced similar effects. (E-N) Dorsal views of embryos that at HH5/6 were treated with control-or Bmp-loaded “white” beads as indicated at the top of the panel. After 6 hours, the embryos were analysed for the expression of *Isl1* and the caudal PHM-and head skeletal muscle precursors marker *Tbx1*. All Bmp molecules upregulated *Isl1* and suppressed *Tbx1*.

Given that different Bmp molecules may elicit different responses (Klumpe et al., 2022), we tested whether the Bmps associated with heart and lateral mesoderm development, namely the ohnologues Bmp2 and Bmp4 and the more distantly related paralogues Bmp7 and 10 (Hinck et al., 2016; Huminiecki et al., 2009) may all upregulate *Isl1* and downregulate the PHM marker *Tbx1* as previously shown for Bmp2 (Bothe et al., 2011). Indeed, at 6.41 µM, all Bmp ligands induced ectopic *Isl1* expression (Fig.1F-I; green arrowheads) and suppressed *Tbx1* (Fig.1K-N; red arrowheads), whereas control beads had no effect (Fig1.E,J; blue arrowheads).

To explore the time frame in which we can expect robust responses to signalling molecules, we carried out a time course, exposing HH5/6 and HH7/8 embryos to beads with 6.41 µM Bmp2 (Fig.2A- R) or 5.35-10.7 μM Wnt3a (Fig.2S-X; data shown for HH7/8 and 8.56 μM Wnt3a) for 2 hours, 4 hours and 6 hours. The direct Bmp target gene *Msx2* (Sirard et al., 2000), began to respond after 4 hours at HH5/6 and 2 hours at HH7/8; responses were robust after 6 hours for both stages (Fig,2A-C, M-O; green arrowheads). *Isl1* showed a weak response after 2 hours at HH5/6, a more pronounced response after 2 hours at HH7/8, and readily detectable responses after 6 hours for both stages (Fig.2D-F, P-R; green arrowheads). Similarly, at HH5/6 the cardiac and lateral mesoderm marker *Hand2* was mildly upregulated within 4 hours, robustly within 6 hours (Fig.2G-I; green arrowheads); *Tbx1* however was already downregulated within 2 hours at this stage (Fg.2J-L; red arrowheads). When testing the direct Wnt target *Lef1* (Li et al., 2006) at HH7/8, we found expression was upregulated and rostrally expanded from its somitic domain within 2 hours (Fig.2S-U, green arrowheads). Further, *Cyp26C1*, a PHM marker and retinoic acid inhibitor we identified here as a highly Wnt-sensitive gene (see section 3.6), was slightly downregulated within 2 hours, robustly within 6 hours (Fig.2V-X, red arrowheads). We thus concluded that a 6 hours treatment regime will allow to detect robust responses of direct and first-wave indirect target genes, and will reveal which developmental options are available to the PHM.

**Figure 2.**
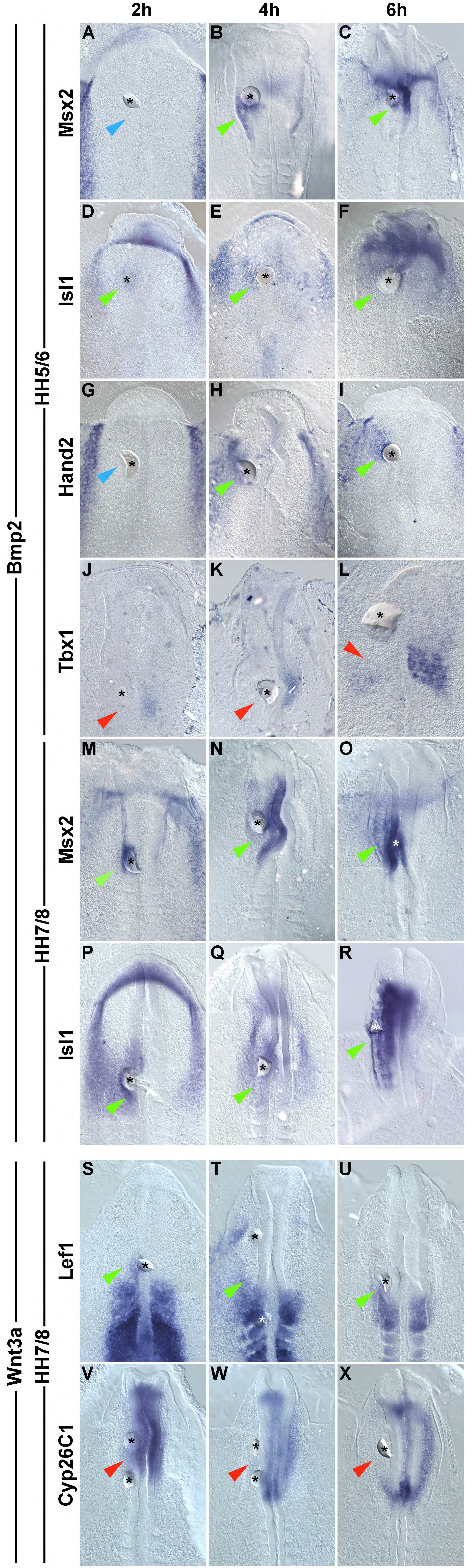
Time-course to establish the onset of Bmp and Wnt effects on the PHM. Dorsal views of embryos that had received white beads loaded with 6.41µM Bmp2 (A-R) or 8.56 µM Wnt3a (S-X) at stage HH5/6 (A-L) or HH7/8 (M-X) as indicated on the left of the panel. Embryos were re-incubated for 2 hours, 4 hours or 6 hours as indicated at the top of the panel. Thereafter, the embryos were assayed for the expression of genes known to be upregulated by Bmp (*Msx2, Isl1, Hand2*), downregulated by Bmp (*Tbx1*) and upregulated by β-Catenin-mediated Wnt signalling (*Lef1*). We also included *Cyp26C1*, a gene we found to be readily suppressed by Wnt3a (see also Fig.14). We found that upregulation (green arrowheads) and downregulation (red arrowheads) became detectable in 2-4 hours; robust changes of expression typically required 6 hours of exposure.

#### 3.1.2. Test for the PHM’s responsiveness to Bmp and Wnt from early neurula to early organogenesis stages

Having established that Bmp and Wnt ligands delivered on beads reliably affect gene expression within 6 hours, we now tested whether within this 6-hour time frame the PHM at stages HH5/6, HH7/8, HH9/10 and HH13/14 would be responsive to the ligands. These stages were selected because of earlier observations on stage-specific expression patterns and responses to signalling molecules: at HH5/6, genes driving cardiac development and the first PHM genes are expressed, at HH7/8 the next set of PHM genes becomes activated, at HH9/10, specifically *Msc* (= *MyoR*), a gene controlling the activation of *Myf5* in head skeletal muscle precursors is activated, and at HH13/14 head skeletal muscle differentiation commences ((Bothe et al., 2011; Bothe and Dietrich, 2006; Meireles Nogueira et al., 2015; Moncaut et al., 2012); summarised in Supplementary Figure SF1)). For our first test, we focused on *Msx2* as a direct Bmp target (Sirard et al., 2000) and *Lef1* as a direct Wnt/β-Catenin target gene (Li et al., 2006). *Msx2* is initially expressed in the early lateral mesoderm and the neural folds/ dorsal neural tube; later expression occurs at many sites including the distal aspect of the pharyngeal arches (http://geisha.arizona.edu/geisha/), where the gene contributes to the survival of SHF cells (Chen et al., 2007). We found that Bmp2 strongly activated *Msx2* transcription in the head mesoderm, the neural tube and the surface ectoderm at all stages (Fig3.A-D, green arrowheads, and cross sections Fig.3Bsi-iii). Upregulation also occurred in the HH13/14 neural crest cells and the somites (Fig.3Dii,iii, green arrowheads). *Lef1* is initially expressed in the developing somites and in the caudal aspect of the PHM; later expression domains include the pharyngeal arches, cranial ganglia and the optic cup (http://geisha.arizona.edu/geisha/). At HH5/6 through to HH9/10, Wnt3a rostrally expanded the expression domain of *Lef1* in the caudal PHM (Fig.3E-G, green arrowheads); this effect was specific to the mesoderm (Fig.3Fsi-ii). At HH13/14, Wnt3a upregulated *Lef1* expression in the pharyngeal arches, the neural tube and in the trunk ectoderm (Fig.3H, green arrowheads). Taken together, this suggests that the PHM is sensitive to Bmp and Wnt throughout the stages under investigation.

**Figure 3.**
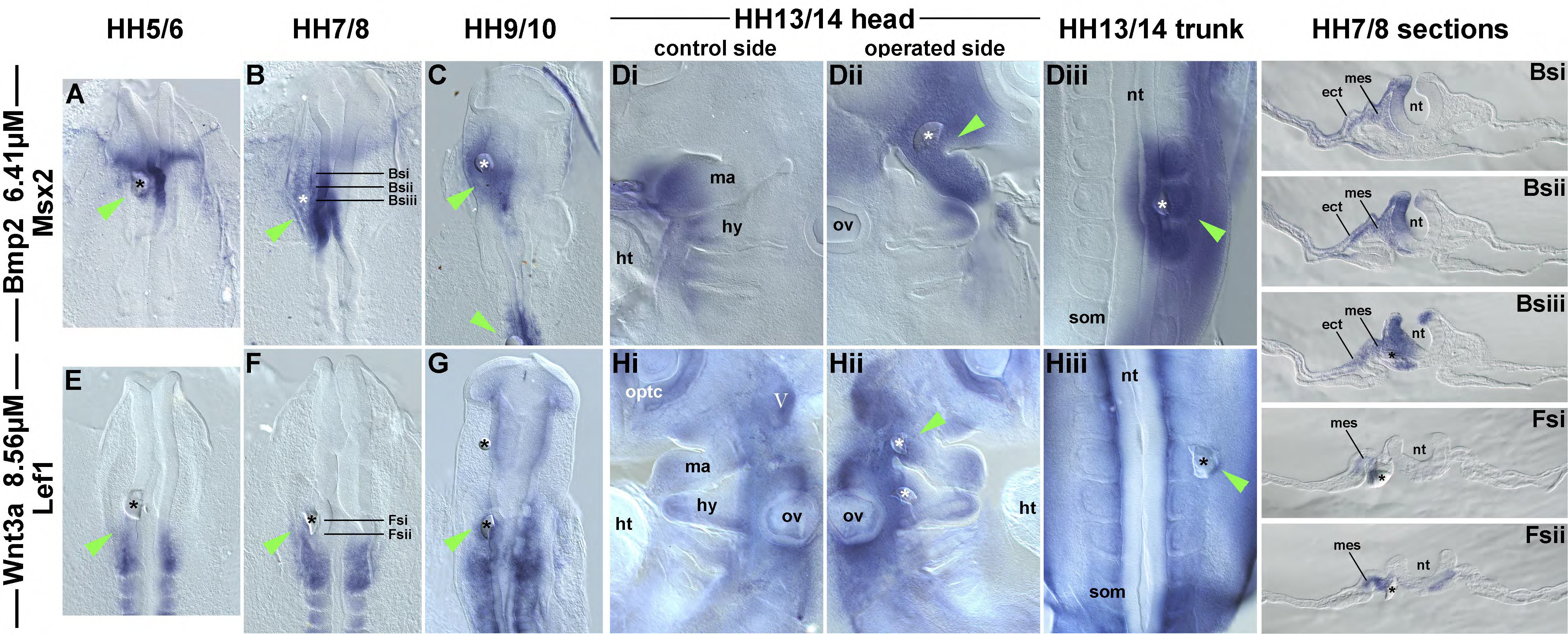
The PHM is Bmp-and Wnt-responsive throughout neurula and early organogenesis stages. Dorsal views (A-C, Diii; E-G; Giii), lateral views of mid-sagittally split specimen (Di-ii; Hi-ii) and cross sections (Bsi-iii; Fsi-ii) of embryos that were treated from early neurula stages at HH5/6 to early organogenesis stages at HH13/14 as indicated at the top of the panel. As indicated on the left, the treatment was with Bmp2 (A,B,C,Dii,Diii, sections Bsi-iii) or Wnt3a (E,F,G,Hii,Hiii, sections Fsi-ii) delivered on white beads. The beads were implanted into the PHM (A-C, Dii; E-G; Hii; sections) or as a control into the trunk paraxial mesoderm (somites; Diii, Hiii). Embryos were re-incubated for 6 hours and then analysed for the expression of the direct Bmp target *Msx2* and the direct Wnt target *Lef1* as indicated on the left of the panel. Images Di-iii and Hi-iii, respectively, are from the same embryo, with Di, Hi showing the untreated control side. The sections are from the embryos in B,F. The position of the beads is marked by an asterisk; for the whole mounts, upregulated marker gene expression is indicated by a green arrowhead. At all stages, Bmp2 strongly activated *Msx2* in the mesoderm and in the neural and surface ectoderm. Wnt3a-beads upregulated *Lef1* specifically in the mesoderm. Note that at stages HH5/6, HH7/8 and HH9/10, the upregulated *Lef1* expression is a rostral extension of the normal expression domain, indicating that Wnt3a caudalised the tissue. Abbreviations: ect, ectoderm; ht, heart; hy, hyoid (2 ^nd^ pharyngeal) arch; ma, mandibular (1 ^st^ pharyngeal) arch; mes, mesoderm; nt, neural tube; ov; otic vesicle; som, somite.

### 3.2. Response of heart-promoting transcription factors to Bmp-treatment at different stages of development

As the PHM is sensitive to Bmp at all the stages investigated, we now were in the position to ask: will Bmp induce the cardiac programme at all stages? Cardiogenesis is initiated by a combination of transcription factors including Isl1, Nkx2.5, Tbx2, Tbx5, Hand2 and Gata4. Mesodermal cells expressing the pioneer factor Isl1 are thought to still have non-cardiogenic options, however the expression of Nkx2.5 drives cells towards a cardiomyocyte fate (Gao et al., 2019; Jia et al., 2018). Importantly, Nkx2.5 is not sufficient for cardiomyocyte differentiation, and it is the combinatorial expression of cardiogenic transcription factors that allows beating cardiomyocytes to form, both in vivo and in vitro (Ieda et al., 2010; Luna-Zurita et al., 2016; Zhou et al., 2012). We therefore included all these markers in our analysis. It has to be considered, however, that their expression patterns are overlapping but not identical: in wildtype embryos, all are expressed in the splanchnic lateral mesoderm that delivers the primitive heart and thus they are associated with cardiogenic precursor cells. *Isl1, Nkx2.5, Tbx2* and *Hand2* expression is maintained in the distal pharyngeal arches that contain the SHF; *Gata4* and *Tbx5* expression are maintained in the caudal SHF; *Tbx5* expression eventually becomes confined to the inflow tract and atria; *Isl1* is switched off in differentiating cells. Notably, *Isl1*, *Nkx2.5* and *Tbx2* are also expressed in the pharyngeal endoderm; *Isl1* is further expressed in the dorsal neural tube and cranial ganglia; *Hand2, Gata4, Tbx2* and *Tbx5* are more widely expressed, including the lateral and extraembryonic mesoderm of the trunk (http://geisha.arizona.edu/geisha/).

Implantation of Bmp2-beads into the PHM triggered expression of all the cardiac marker genes at HH5/6 (Fig.4A,E,I,M,Q,U; green arrowheads), control beads had no effect (not shown). Upregulation was strong for *Isl1, Nkx2.5, Hand2* and *Gata4*, robust for *Tbx2* and more limited for *Tbx5*, likely because *Tbx5* is an indirect target (see also (Yamada et al., 2000)). Notably, at HH7/8, Bmp2 failed to activate *Tbx5* in the PHM (Fig.4V, blue arrowhead), and from HH9/10 onwards, Bmp2 failed to activate *Gata4* (Fig.4S, blue arrowhead). At HH13/14, Bmp-beads dorsally expanded the expression domains of *Isl1* and *Hand2* which are normally confined to the distal/ventral pharyngeal arches that contain the SHF (Fig.4Dii, Pii, green arrow). However, ectopic expression around the bead was not observed. Moreover, expression of *Nkx2.5*, *Tbx2*, *Gata4*, *Tbx5* was unchanged (Fig.4Hii, Lii, Tii, Xii, blue arrowheads). When Bmp-beads were implanted into the trunk paraxial mesoderm, i.e. the somites, expression of *Tbx2*, *Hand2*, and *Gata4* was upregulated (Fig.4Liii, Piii, Tiii, green arrowheads); the other markers did not respond. The activation of these more general lateral mesoderm markers suggests that Bmp can ‘lateralise/ventralise’ the paraxial mesoderm for some time, both in the head and in the trunk. However, the ability of Bmp to activate the full cardiac marker set fades away after stage HH5/6.

**Figure 4.**
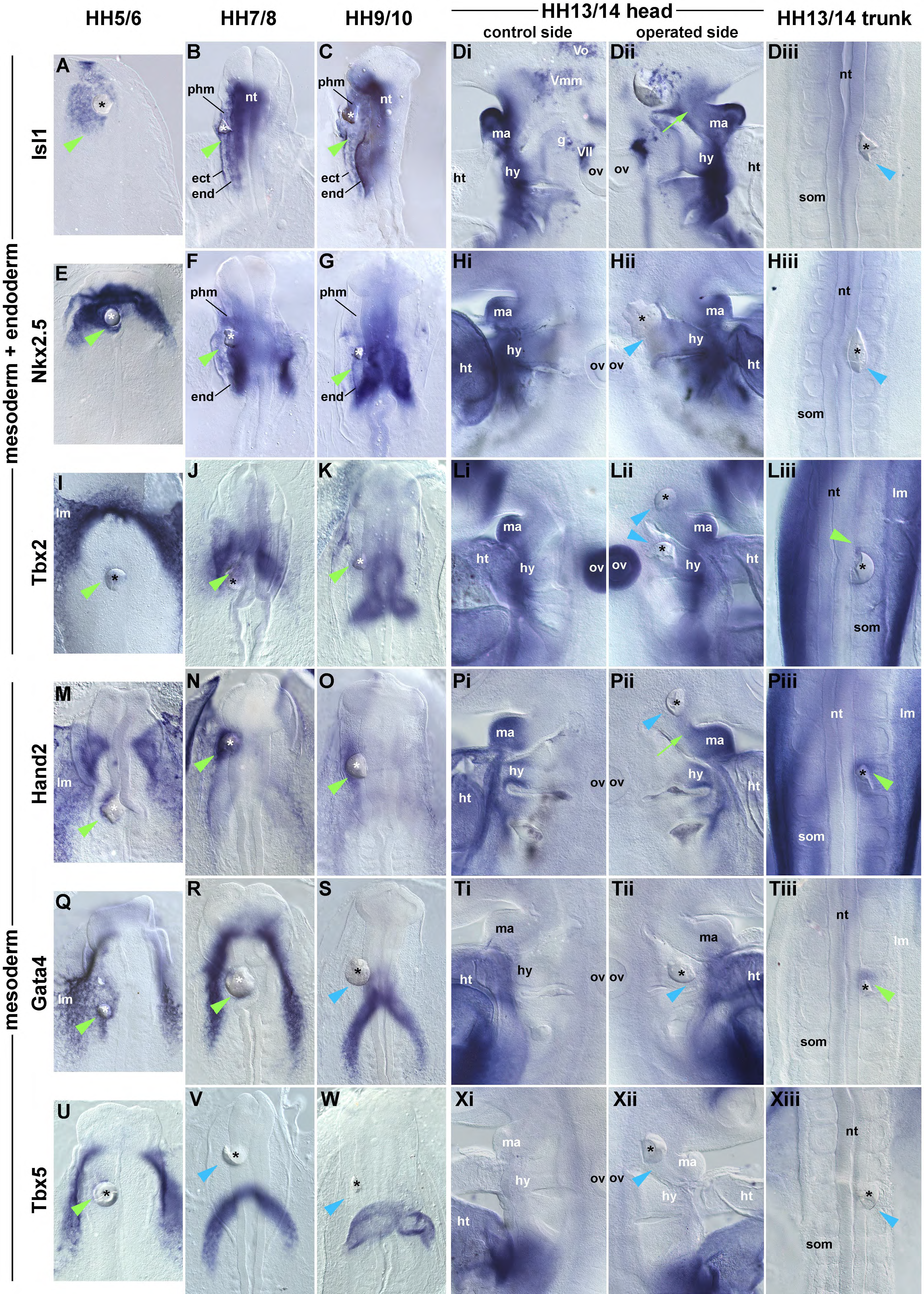
Bmp activates the full set of cardiac markers at HH5/6 only. Beads loaded with Bmp2 were implanted into the PHM or somites of embryos at HH5/6, HH7/8, HH9/10 and HH13/14 as shown in Fig.3 A-Diii. After 6 hours of re-incubation, the embryos were assayed for genes expressed in the lateral/ cardiac mesoderm and endoderm or in the lateral/ cardiac mesoderm alone as indicated on the left. Importantly, these genes cooperatively drive cardiogenesis. The specimen are shown in dorsal or lateral views as in Fig.3; upregulated gene expression around the bead is indicated by green arrowheads, expanded expression domains by green arrows, unchanged expression by blue arrowheads. In the PHM, Bmp activated the full set of marker genes only at HH5/6; in the trunk, generic lateral mesoderm markers were upregulated also at later stages. Abbreviations: ect, surface ectoderm; end, endoderm; g, geniculate ganglion; ht, heart; hy, hyoid (2^nd^ pharyngeal) arch; lm, lateral mesoderm; ma, mandibular (1^st^ pharyngeal) arch; nt, neural tube; ov; otic vesicle; phm, paraxial head mesoderm; som, somite; Vo, ophthalmic division of the trigeminal (5^th^ cranial) ganglion; Vmm maxillomandibular division of the (5^th^ cranial) trigeminal ganglion, VII, facial (7^th^ cranial) ganglion.

### 3.3. Cardiac differentiation of the PHM

#### 3.3.1. Cardiac differentiation of PHM induced by Bmp

The activation of the complete set of cardiac precursor markers at stage HH5/6 suggested that the PHM may have cardiac competence, but only at this early stage. However, using the same 6-hour treatment regime, Bmp did not activate *Myocd*, *Mef2c* and cardiac effector genes, even when higher concentrations were used (Supplementary Figure SF2). Yet in the embryo it takes some 14 hours from stage HH5/6 to HH10 when the first cardiomyocytes are terminally differentiated and the heart begins to beat (reviewed in (Wittig and Münsterberg, 2020)). We thus allowed the beaded embryos to develop for 16-18 hours overnight. Moreover, we trebled the Bmp concentrations to compensate for diffusion away from the bead. Terminal differentiation was revealed by immunofluorescence with the MF20 antibody which recognises sarcomeric myosin (Fig.5, red staining). Notably, at elevated concentrations, Bmp triggered ectopic sarcomeric myosin expression in the PHM, albeit at low frequency (see Supplementary Table ST1, n=4:10). Importantly, sarcomeric myosin expression was only obtained when the bead was inserted at HH5/6 (Fig.5L, green arrowhead; inset: confocal image of the MF20 positive region).

**Figure 5.**
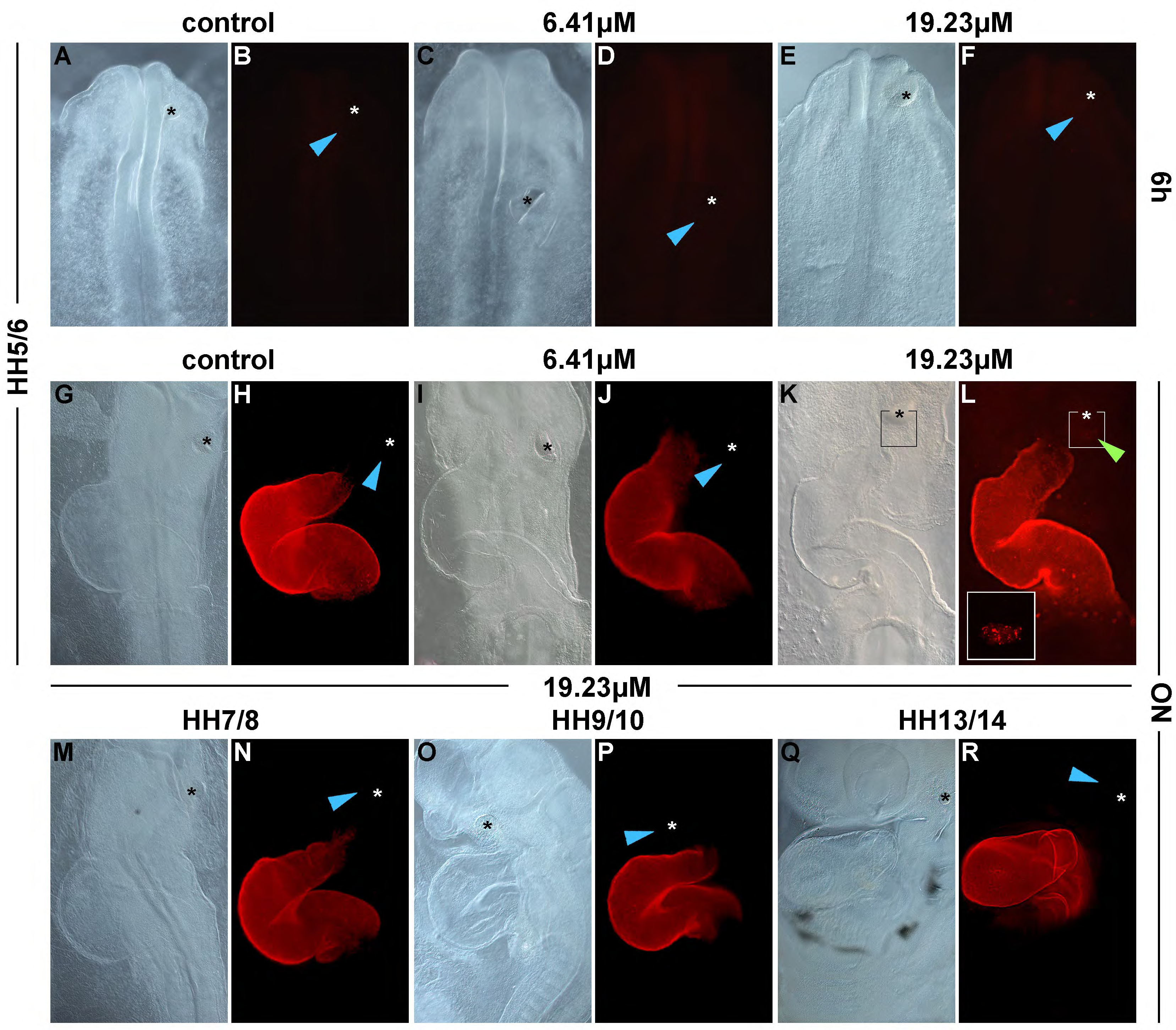
Longer exposure to Bmp allows the HH5/6 PHM to complete myocardial differentiation. Implantation of control beads and beads loaded with 6.41µM or 19.23µM Bmp2 into the HH5/6, HH7/8, HH9/10 and HH13/14 PHM, followed by MF20-immunofluorescence to detect sarcomeric myosin (red staining). Embryos were left to develop for 6 hours (A-F) or overnight (G-R). (A,C,E,G,I,K,M,O,Q) show DIC images, (B,D,F,H,J,L,N,P,R) show the same embryo with fluorescence microscopy, respectively. Images were collected as z-stacks, the heart and the beads in the PHM were compressed into one image to show both in focus. The inset in (L) is a confocal image if of the boxed region in (K,L). (A-F) are dorsal views; (G-R) are ventrolateral views. Ectopic cardiomyocyte differentiation occurred around the bead after overnight incubation with 19.23µM Bmp2, but only in HH5/6 hosts (L, green arrowhead and inset).

#### 3.3.1. Cardiac differentiation of PHM the cardiogenic environment of the embryo

The implantation of Bmp-loaded beads only partially mimics the complement of factors cardiogenic cell are normally exposed to. Moreover, the Bmp concentrations used here may be artificially high. We therefore tested whether the PHM can complete differentiation in the natural cardiogenic environment of the embryo. To achieve this, the PHM was excised from GFP-or TdTomato-expressing embryos and transplanted into the cardiogenic region of unlabelled hosts at HH5/6 and HH7/8. The embryos were allowed to develop over night as before. The MF20 antibody was used to assay for terminal differentiation; GFP or TdTomato expression was enhanced using an anti-GFP or pan-anti-RFP antibody, respectively. The location of grafted cells was established by fluorescence microscopy (Fig.6A-J), co-expression of sarcomeric myosin and GFP or TdTomato was analysed by confocal microscopy (Fig.6i-iii insets). For TdTomato grafts, the red and green channels were inversed such that throughout Fig.6, all grafted cells are displayed in green, all MF20 staining is in red. N-numbers for the grafting experiments are displayed in the Supplementary Table ST2.

**Figure 6.**
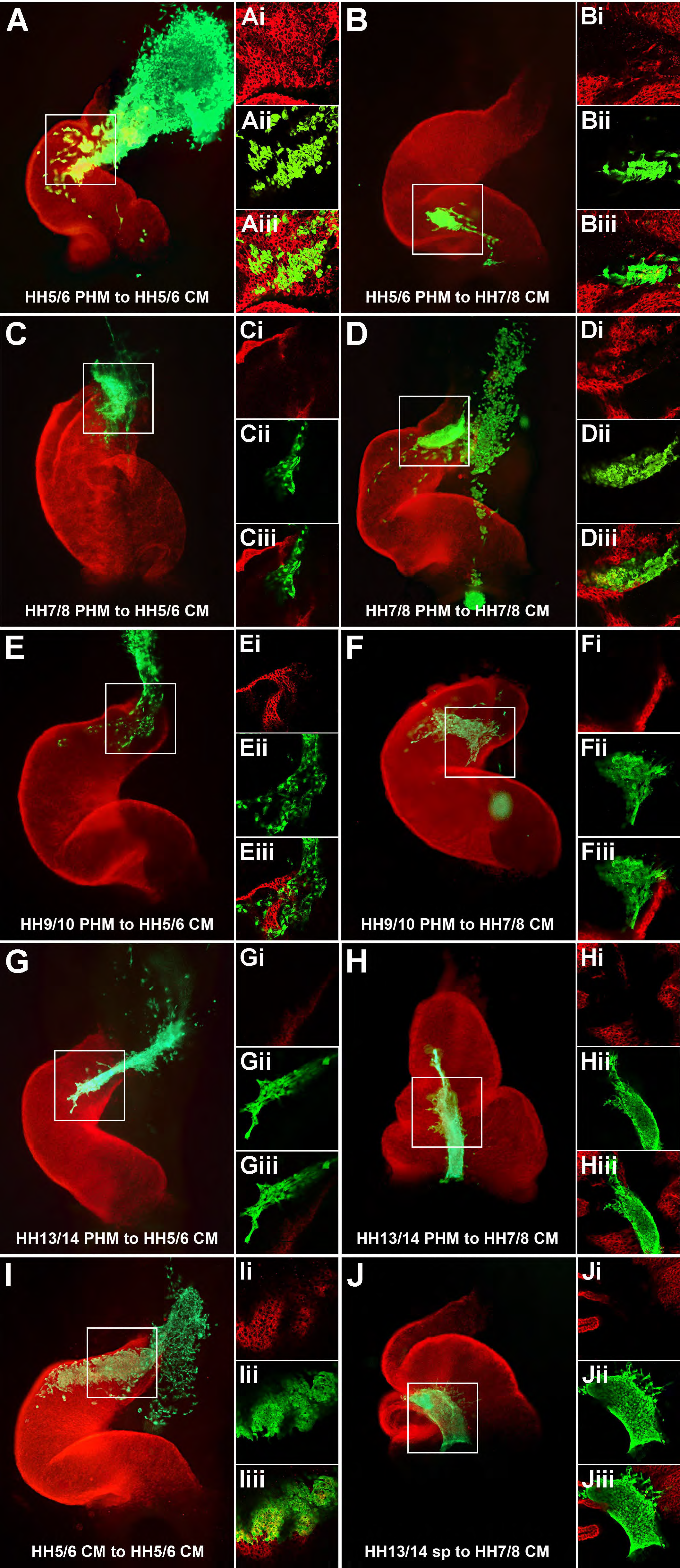
The HH5/6 PHM can complete myocardial differentiation in vivo. PHM (A-H), lateral/ cardiac mesoderm (CM, positive control; I) or trunk segmental plate (sp, negative control; J) from fluorescing donors was implanted into unlabelled hosts at HH5/6 (left panel) or HH7/8 (right panel) as indicated for each image set. After overnight re-incubation, immunofluorescence to detect sarcomeric myosin (red) and to enhance the fluorescence of the graft (green) was performed. (A,B,C,D,E,F,G,H,I,J) are images of the heart obtained by compound fluorescence microscopy, ventral views; (i, ii, iii) are the corresponding confocal sections showing the red channel (i), the green channel (ii) and the overlay (iii). Note that the PHM derived from a HH5/6 donor differentiated when grafted into the cardiogenic environment of a HH5/6 host (A-Aiii), very faint sarcomeric myosin expression was also obtained in a HH7/8 host (B-Biii). Faint MF20-positivity was also observed in HH7/8 grafts albeit rarely (D-Diii). PHM from older stages of development (E-Hiii) or segmental plate grafts (J-Jiii) became incorporated into the host heart but did not differentiate. HH5/6 cardiogenic lateral head mesoderm readily differentiated (I-Iiii). Abbreviations: CM, cardiac mesoderm; PHM, paraxial head mesoderm; sp, segmental plate (pre-somitic mesoderm).

We observed that the retention rate of tissue grafted into the cardiogenic region at HH5/6 was varied, and grafts often dispersed, possibly because of the morphogenetic movements that lengthen the head at this time ((Cui et al., 2009; Redkar et al., 2001); for retention rates, see also ST2). Nonetheless, PHM derived from HH5/6 donors was able to differentiate and express sarcomeric myosin, with larger areas displaying MF20-positivity in HH5/6 compared to HH7/8 hosts (Fig.6A,B; MF20-positivity in n=3:7 for HH5/6 hosts, n=4:12 for HH7/8 hosts). Grafts derived from HH7/8 donors very rarely differentiated (Fig.6C,D; n=0:3 in HH5/6 hosts, 2:11 in HH7/8 hosts). For PHM from HH9/10 donors (Fig.6E,F; n=0:11) or HH13/14 donors (Fig.6G,H; n=0:6), MF20-positivity was not observed at all, despite the fact that these grafts had moved into the heart together with endogenous cardiogenic tissue, and were surrounded by differentiated myocardium. As positive control, we used HH5/6 cardiogenic mesoderm which is specified but not determined to undertake cardiogenesis. This tissue differentiated in n=6:7 HH5/6 hosts (Fig.6I) and n=4:6 HH7/8 hosts (not shown). As negative control we used segmental plate tissue derived from HH13/14 donors, which was transported into the heart but did not differentiate (n=0:3, Fig.6J). Taken together, this suggests that at stage HH5/6, the PHM is specified but not determined to follow paraxial programmes, and it has cardiac competence. This competence fades away from HH7/8 onwards.

### 3.4. Non-myocardial programmes stimulated by Bmp in the PHM after the loss of cardiac competence

Given that Bmp alone was sufficient to trigger the cardiac programme in the HH5/6 PHM, yet the PHM was Bmp-sensitive at all stages tested here, we wondered which programmes Bmp may control after the loss of cardiac competence. To test this, we took two approaches. (1) We performed an RNAseq screen for BMP-dependent genes at HH7-8; for technical reasons (no decay of cells and tissues, sufficient RNA for analysis) all head tissue was used. (2) We performed a candidate gene approach, testing whether and at which stages markers for other programmes available to the head mesoderm may be regulated by Bmp.

#### 3.4.1. RNAseq screen for Bmp-dependent genes at HH7-8

To identify Bmp-dependent genes, we exposed embryos at HH7-8 to the ALK2/3 inhibitor LDN193189 for 6 hours, and analysed the downregulation of genes in the head tissue by RNAseq. Notably, genes downregulated after inhibition of Bmp signalling included genes like *Noggin*, *Msx2*, *Isl1*, *Nkx2.5*, *Tbx2*, and *Gata4* (Fig.7A) which were experimentally induced by Bmp at this stage (see Fig.3 and ST2). We used these genes to define our thresholds for a significant downregulation. 98 protein-coding genes fulfilled our criteria for inclusion (Fig.7A and Supplementary Table ST3). Interestingly, the most downregulated gene was the chicken orthologue of the Wnt inhibitor *Szl*. Among the downregulated, Bmp-dependent genes were also the known Bmp targets *Id1-4*.

**Figure 7.**
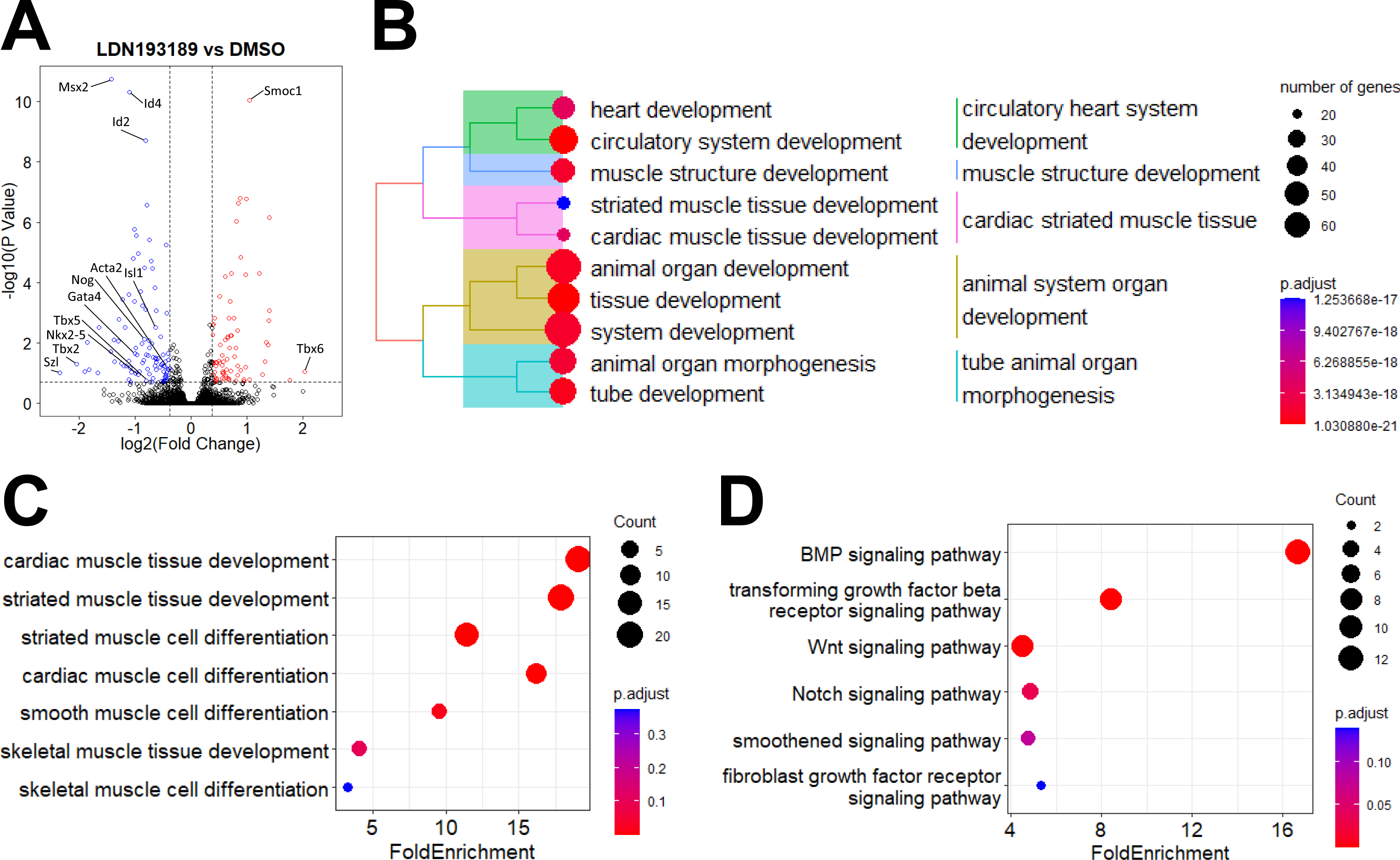
RNASeq analysis of Bmp-dependent genes in the early head. HH7-8 embryos were cultured for 6 hours in culture medium with 1:1000 DMSO (control) or 1μM LDN193189. RNA was prepared from the head region to the first somite, and analysed by RNASeq. (A) Volcano plot of the 17,007 protein-coding genes that were mapped. Marked are the thresholds for the p-adjusted and fold change parameters. The 98 genes downregulated (blue) and 78 genes upregulated (red) after LDN193189 treatment are indicated. Marked are the top up-and downregulated genes by p-adjusted value and fold change, and the genes found to be activated by Bmp in this study. (B) Treeplot (hierarchical clustering) of the top 10 ‘Biological Process’ GO terms, based on over-representation analysis of the genes downregulated by LDN19318 treatment. The list is dominated by GO terms associated with cardiac muscle development. The remaining GO terms all relate to embryonic development, indicating the high representation of developmental control genes in the gene set. (C) Dotplot to illustrate the representation of ‘Biological Process’ GO terms associated with the development of different muscle types in the set of genes downregulated by LDN19318 treatment. GO terms are sorted by p-adjusted value. Cardiac/striated muscle and smooth muscle are over-represented. (D) Dotplot to illustrate the representation of ‘Biological Process’ GO terms associated with major signalling pathways in the set of genes downregulated by LDN19318 treatment. GO terms are sorted by p-adjusted value. BMP, TGFβ, Wnt, and Notch signalling are over-represented, while Fgf and hedgehog/smoothened signalling are below the threshold of 0.05 false detection rate. In B-D, circle size is relative to the number of genes annotated for the respective GO term, while the colour represents the p-adjusted value (as defined in the respective keys).

Overrepresentation analysis for ‘Biological Process’ GO terms indicated that many of the Bmp-dependent genes are known to be involved in embryonic development, and in particular highlighted GO terms linked with cardiac muscle formation (Fig.7B). This was expected, given the dominance of the cardiac programme in the head at this stage. Since stage HH7/8 coincides with the diminishing cardiac competence in the PHM, we tested the gene set for alternative mesodermal programmes by investigating the representation of GO terms associated with different muscle types (Fig.7C). Cardiac muscle GO terms, largely overlapping with striated muscle GO terms were highly represented in our gene set, but we also found overrepresentation of the smooth muscle differentiation GO term featuring genes such as *Acta2* and *Csrp2*. The GO terms for skeletal muscle are populated by genes involved in trunk skeletal muscle rather than the distinct head skeletal muscle programme, and hence are not informative.

To investigate if Bmp signalling could be linked with the regulation of other signalling pathways, we interrogated our gene set for overrepresentation of the ‘Biological Process’ GO terms for major signalling pathways (Fig.7D). As expected, the GO terms for Bmp signalling and the sister Tgfβ pathway were highly represented. However, we also found the Wnt signalling GO term with genes like *Szl* and *Dact1* overrepresented, suggesting a link between Bmp and Wnt signalling. Also overrepresented was the Notch-signalling GO term.

#### 3.4.2. Bmp effect on candidate markers associated with non-myocardial programmes

##### 3.4.2.1. Bmp effects on vascular endothelial, endocardial, smooth muscle and endo-ectodermal markers

The head mesoderm is thought to deliver the craniofacial vasculature (Couly et al., 1995; Noden, 1989), and in the trunk, vascular endothelial cell development from paraxial mesoderm (somites) depends on Bmp (Nimmagadda et al., 2005). Moreover, the lateral head mesoderm, besides forming the myocardium, also delivers the specialist vascular endothelial cells of the endocardium (Harris and Black, 2010; Misfeldt et al., 2009; Sendra et al., 2021). We thus returned to the 6-hours treatment of the PHM with Bmp beads, this time assaying for the vascular endothelial precursor (angioblast) marker *Kdr,* also known as *Flk1* or *VegfR2*, and the endocardial marker *NfatC1* (Nimmagadda et al., 2005; Wu et al., 2011). The latter is also expressed in the pharyngeal endoderm. As expected, Bmp stimulated the expression of *Kdr* in the somites (Fig.8Diii, green arrowhead). Yet between stages HH5-10, Bmp reduced *Kdr* expression in the head (Fig.8A-C, red arrowheads), which warrants further investigation. Notably, Kdr expression was not affected by LDN193189 in our RNAseq screen. The expression of *NfatC1* was not changed in the mesoderm, but at HH13/14, expression was upregulated inside the embryo in the pharyngeal endoderm (Fig.8Hii, green arrows). Taken together, Bmp seems not to shift head mesodermal cells towards a vascular endothelial fate.

**Figure 8.**
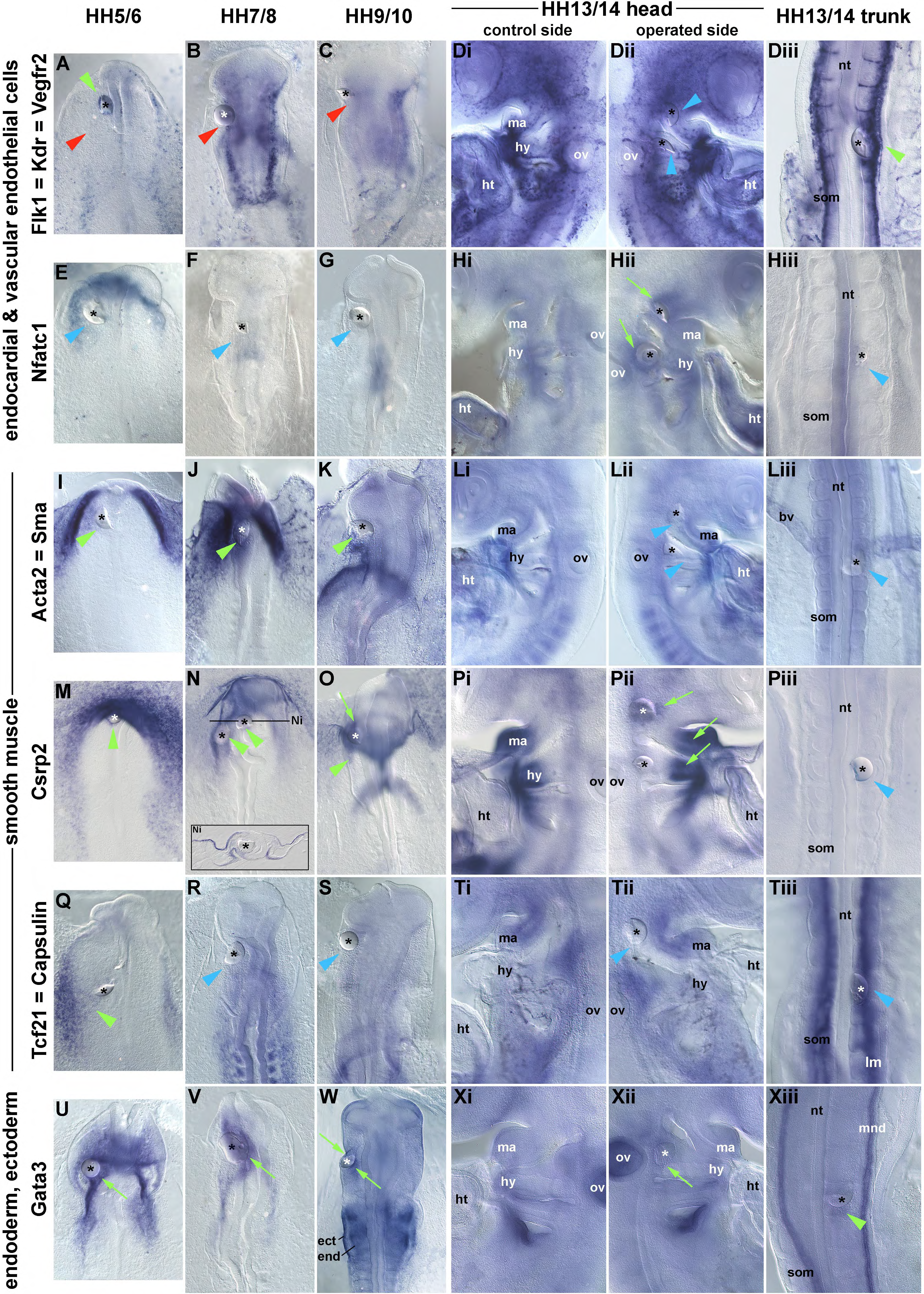
Responses to Bmp by markers associated with vascular endothelial and smooth muscle fates and with endoderm/surface ectoderm patterning. Dorsal or lateral views of embryos that had received Bmp beads into the PHM as in Fig.3,4; duration of treatment was 6 hours. Green arrowheads indicate upregulated, red arrowheads downregulated, blue arrowheads wildtype expression in the PHM or in the somite; green arrows indicate responses in other expression domains. Abbreviations as before and: ect, surface ectoderm; end, endoderm; mnd, mesonephric duct. *Flk1*, also known as *Kdr* or *Vegfr2*, labels developing vascular endothelial cells. Besides an occasional accumulation of Kdr+ cells around the bead (A, green arrowhead), *Kdr* was downregulated in the head between HH5-10 (A-C, red arrowheads), but upregulated in the trunk at HH13/14 (Diii, green arrowhead). *NfatC1* labels the vascular endothelial cells forming the endocardium. However, the gene is also expressed in the pharyngeal endoderm; this expression was upregulated at HH13/14 (Hii, green arrows). *Acta2 (Sma)*, is initially expressed widely in the cardiogenic and head lateral mesoderm. Expression occurs temporarily also in skeletal muscle. However, prolonged gene expression and function distinguishes smooth muscle. The gene was upregulated by Bmp from HH5- HH10(I-K,greenarrowheads). *Csrp2*, a gene controlling smooth muscle development but also expressed in the lateral ectoderm and endoderm was upregulated throughout HH5-10; the section indicates that this upregulation encompassed the mesoderm (Ni). At HH13/14, Bmp slightly expanded the expression domain in the distal pharyngeal arches and increased expression in the trigeminal placode (Pii, green arrows). *Tcf21* (*Capsulin*), a gene widely expressed in intermediate and lateral mesoderm derivatives, promotes fibroblast and suppresses smooth muscle formation in the epicardium. The gene was mildly upregulated at HH5/6 (Q, green arrowhead). *Gata3* labels the endoderm and surface ectoderm, and at later stages, the mesonephric duct (mnd). In the head Gata3 was upregulated by Bmp in the ecto-and endoderm at all stages (U-Xii, green arrows, in the trunk expression was upregulated in the somite green arrowhead).

The lateral mesoderm delivers the smooth muscle collar to complete the cardiac outflow tract (Waldo et al., 2005; Wu et al., 2006), reviewed by (Donadon and Santoro, 2021), and we thus reasoned that Bmp may promote the smooth muscle programme. Markers specific for smooth muscle are limited. However, *Acta2* (*Sma*), whilst expressed initially in all developing muscle, is only retained in smooth muscle, and *Csrp2* promotes smooth muscle development (Chang et al., 2003; Lopez-Sanchez et al., 2009), reviewed by (Donadon and Santoro, 2021); both genes were identified as Bmp-dependent in our RNAseq screen. *Tcf21* (*Capsulin*) is widely expressed in lateral and intermediate mesoderm derivatives and is associated with smooth muscle precursors. However, it suppresses smooth muscle differentiation and in the heart promotes cardiac fibroblast formation from the epicardium (Tandon et al., 2012; von Scheven et al., 2006b), reviewed in (Hu et al., 2020). We found that between HH5-10, Bmp strongly upregulated *Acta2* in the PHM (Fig.8I-K, green arrowheads); at HH13/14, *Acta2* did not respond any longer (Fig.8Li-iii). At stages HH5-10, *Csrp2* is predominantly expressed in the lateral ectoderm and the endoderm underlying the cardiac precursors. Yet Bmp upregulated *Csrp2* in all three germ layers including the mesoderm (Fig.8M-O and section Ni; green arrowheads). At HH13/14, the response was limited as Bmp slightly expanded the *Csrp2* expression domain in the distal pharyngeal arches and elevated expression levels in the trigeminal placode (Fig.8Pii, green arrowheads). The expression domain of *Tcf21* somewhat expanded towards a Bmp-loaded bead placed into the PHM at HH5/6 (Fig.8Q, green arrowhead), and did not change at later stages (Fig.8R-Tiii, blue arrowheads) neither in the head nor the trunk. Together this suggests that smooth muscle competence is retained in the PHM longer than cardiac competence.

Since in the embryo, signals to influence cardiac development to a large extent originate from the pharyngeal endoderm and encompass more than Bmp (Warkman et al., 2008), because the endoderm of the anterior intestinal portal has been suggested as a cardiac organiser (Anderson et al., 2016), and because *NfatC1* and *Csrp2* were upregulated by Bmp in the endoderm, we had a further look at this tissue. Using *Gata3* as a marker for the cranial endoderm and ectoderm and in the trunk, the mesonephric duct (Bothe and Dietrich, 2006; Sheng and Stern, 1999), we found that this marker was mildly upregulated in the ectoderm and endoderm at all stages (Fig.8U-Xii, green arrows) and also in the trunk paraxial mesoderm (somite) at stage HH13/14 (Fig.8Xiii; green arrowhead). This suggests that the loss of cardiac competence in the PHM from HH7/8 onwards likely is not due to the endoderm having become refractory to Bmp and thus not producing heart-inducing signals.

##### 3.4.2.2. Bmp effect on head skeletal muscle markers

The PHM is a prominent source of craniofacial skeletal/voluntary muscle, and our previous work suggested that Bmp is involved in the activation of *Msc* (*MyoR*) which later upregulates *Myf5* and *MyoD* ((Bothe et al., 2011; Bothe and Dietrich, 2006; Moncaut et al., 2012), reviewed in (Schubert et al., 2018)). On the other hand, Bmp signalling suppresses skeletal muscle differentiation, both in the head and in the trunk (Tzahor et al., 2003; von Scheven et al., 2006a). To explore this problem, we turned to the three genes that demarcate head skeletal muscle precursors, *Pitx2*, *Tbx1* and *Msc* (Fig.9) and markers associated with entry into head and trunk skeletal muscle differentiation (*Six1*, *Eya1*, *Dach1*, *Dach2*, *Ebf2*, *Myf5*; Fig.10). Since *Pitx2* is expressed in the left lateral mesoderm (reviewed in (Franco et al., 2017)), we investigated both the left and the right PHM of the embryo.

**Figure 9.**
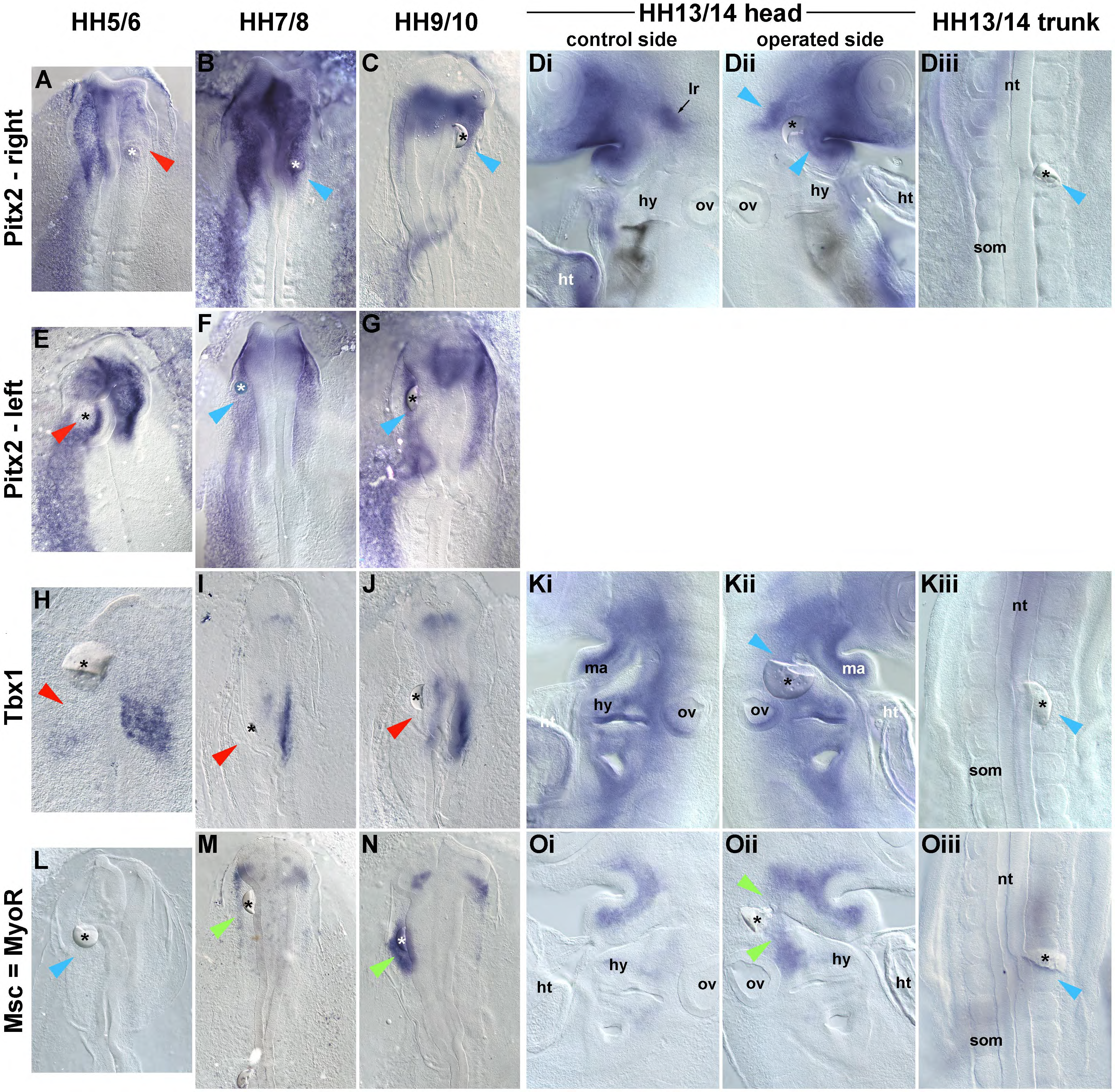
Dynamic responses to Bmp for the upstream regulators of the head skeletal muscle programme. Implantation of Bmp2-beads into the PHM, dorsal or lateral views of embryos and abbreviations as in Fig.3,4,8. After 6h, the embryos were assayed for the expression the three upstream regulators of head skeletal myogenesis, *Pitx2*(A-G), *Tbx1* (H-Kiii) and *Msc*(L-Oiii).Owingtotheasymmetric expression of *Pitx2*, both the right (A-C) and the left side (E-G), were tested at the three neurula stages. Upregulated gene expression around the bead is indicated by green arrowheads, downregulated expression by red arrowheads, unchanged expression by blue arrowheads. At HH5/6, Bmp downregulated *Pitx2* in the PHM, both on the left and the right side (A,E, red arrowheads). The gene became refractory to Bmp from HH7/8 onwards (B-Dii, F-G, blue arrowheads). *Tbx1* was downregulated at HH5-10 (H-J, red arrowheads) and was refractory thereafter. Bmp upregulated *Msc* from HH7/8 onwards (I, J, green arrowheads). This effect was still pronounced at HH13/14 (Kii, green arrowheads).

**Figure 10.**
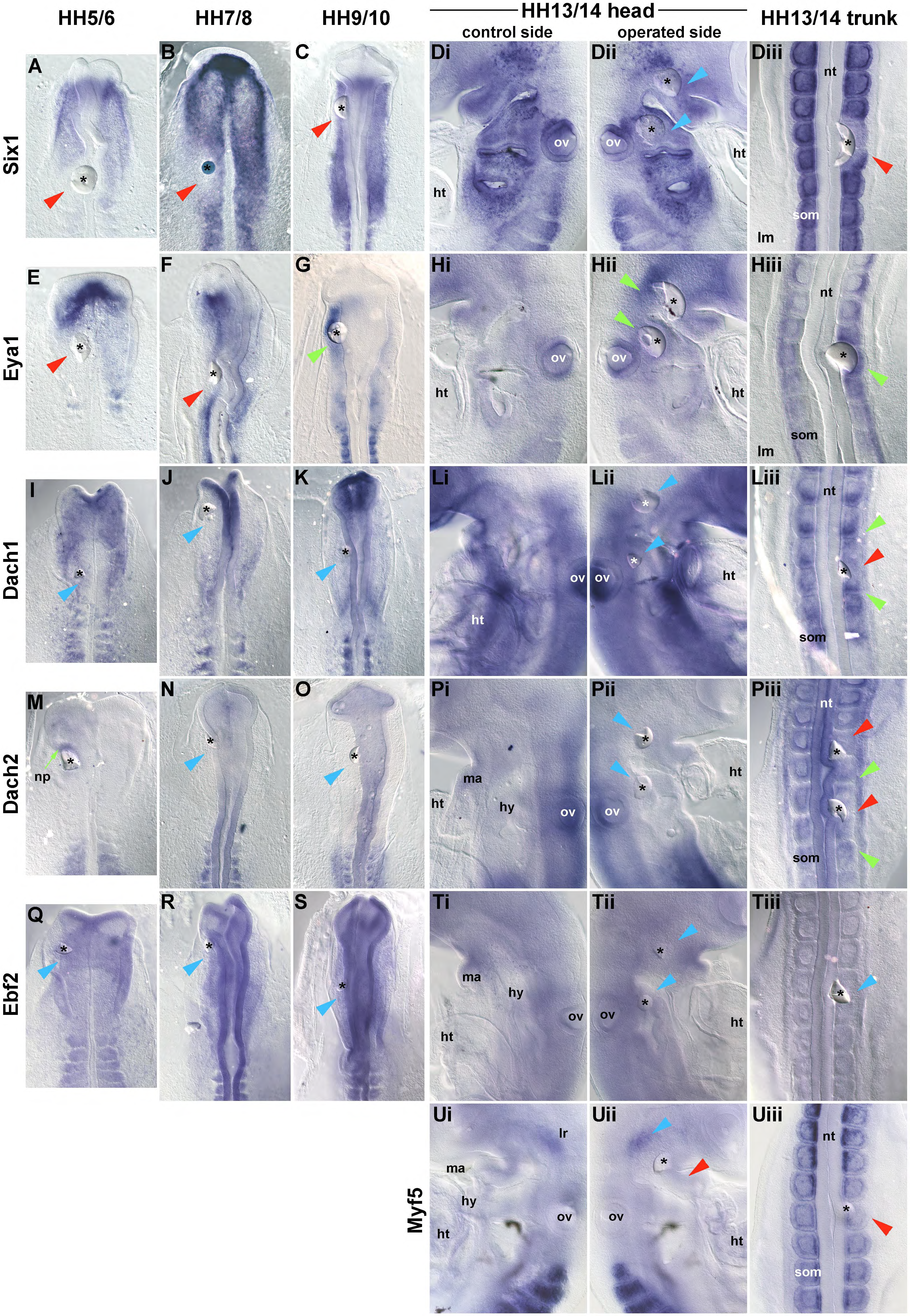
Dynamic responses to Bmp for generic regulators of skeletal muscle formation. Six-hour implantation of Bmp2-beads and annotations and abbreviations as in Fig.3,4,8,9; embryos are shown dorsal or lateral views as before. The green arrow in (M) points at an upregulated expression in the neural plate. Note that initially, Bmp suppressed *Six1* and *Eya2* in the PHM, but this effect faded over time and at HH9/10 and HH13/14, Bmp upregulated *Eya1*, both in the head and in the trunk (G, Hii, Hiii). Bmp did not affect the expression of *Dach1*, *Dach2* and *Ebf2* in the head (I-Lii M-Pii, Q-Tii; blue arrowheads) except an upregulation of *Dach2* in the HH5/6 neural plate (M, green arrow). In the trunk, the Bmp concentration used here downregulated *Dach1* and *Dach2* expression close-by, but upregulated it at a distance (Liii, Piii); *Six1* and *Myf5* were always downregulated (Diii, Uiii; red arrowheads). In the head, *Myf5* was slightly downregulated in the pharyngeal arch muscle anlagen and unaffected in the anlage of the lateral rectus eye muscle (Ui, Uii; lr).

We found that in the PHM, Bmp suppressed *Pitx2* both in the right and left PHM, but only at HH5/6 (Fig.9A,E, red arrowheads). Thereafter, *Pitx2*wasrefractorytoBmp(Fig.9B-Dii,F-G,blue arrowheads). *Tbx1* was downregulated by Bmp at HH5/6, 7/8 and 9/10, in line with the idea that Bmp can lateralise the PHM at these stages (Fig.9H-J, red arrowheads). Thereafter, also *Tbx1* was unaffected (Fig.9Ki-iii). Notably, Bmp began to activate *Msc* at HH7/8 (Fig.9M, green arrowhead), one stage earlier than reported by Bothe (2011), possibly because of the slightly higher Bmp concentration used in the current study. The upregulation of *Msc* was more pronounced at HH9/10 and HH13/14 (Fig.9N, Oii). Yet in the trunk, Bmp did not upregulate *Msc*, indicating a very specific role of Bmp in the activation of the head skeletal muscle programme.

For the genes associated with skeletal muscle differentiation, we found that Bmp downregulated *Six1* at HH5/6 to HH9/10 (Fig.10A-C, red arrowheads); the effect was mild at HH9/10, and no effect wasobservedlater(Fig.10Dii, bluearrowheads). *Eya1* was initially also suppressed (Fig.10E,F; red arrowheads). However, at HH9/10 and 13/14, *Eya1* was upregulated (Fig.10G, Hii; green arrowheads). The expression of the *Dach* and *Ebf* genes was not affected except for some upregulation of *Dach2* in the neural plate (Fig.10M, green arrow). At HH13/14, *Myf5* expression was slightly downregulated in the pharyngeal muscle anlagen (Fig.10Uii, red arrowhead), but not in the anlage of the lateral rectus muscle (Fig.10Uii, blue arrowhead). In the trunk, *Six1* and *Myf5* were downregulated (Fig.10Diii, Uiii), *Dach1* and *Dach2* were downregulated close to the bead and upregulated at a distance where Bmp concentrations are lower (Fig.10Liii, Piii), and *Eya1* was upregulated (Fig.10Hiii). Thus, both in the head and in the trunk, Bmp supports *Eya1* expression, but hinders genes readying cells for terminal skeletal muscle differentiation such as *Myf5*. Taken together, our data specifically on *Msc* and *Eya1* suggest that at the time that cardiac competence of the PHM fades, Bmp promotes the entry into the programme for head skeletal myogenesis.

### 3.5. Role of Wnt/β-Catenin signalling in the control of the cardiac versus the head skeletal muscle programme

Our study revealed that the PHM loses cardiac competence concomitant with the establishment of head skeletal muscle competence. Previous studies inferred that levels of Wnt/β-Catenin signalling might rise in the PHM at stages when we found the PHM’s cardiac competence fades (Tzahor and Lassar, 2001). Thus, Wnt might promote the switch from cardiac to skeletal muscle competence observed here. On the other hand, Wnt/ β-Catenin signalling has been shown to suppress skeletal muscle formation in the head ((Tzahor et al., 2003) and Supplementary Fig. SF1). Wnt however has also been shown to contribute to the caudalisation of the embryo and hence may have a more generic role in suppressing rostral programmes (Marvin et al., 2001). To solve this conundrum, we used the 6-hours experimental paradigm and performed three further tests: (1) We determined when exactly cardiogenesis is suppressed by Wnt3a, a ligand known to trigger the β-Catenin pathway ((Tzahor et al., 2003); Fig.11). (2) We turned to Szl= LOC417741, a Crescent-and Sfrp1/2/5- related Wnt inhibitor strongly expressed in the early cardiac mesoderm and the SHF (Wittler et al., 2008) and the top-ranked Bmp-dependent gene identified in our RNAseq screen, to find out whether Bmp would induce *Szl* in the PHM only at the early, cardiac-competent stage of development (Fig.12A-Dii), and whether Wnt3a would suppress *Szl* similar to the cardiac markers tested before (Fig.12E-Hii). (3) We tested whether Wnt3a would accelerate or repress the markers for head skeletal muscle precursors and the onset of skeletal muscle differentiation (Fig.13).

**Figure 11.**
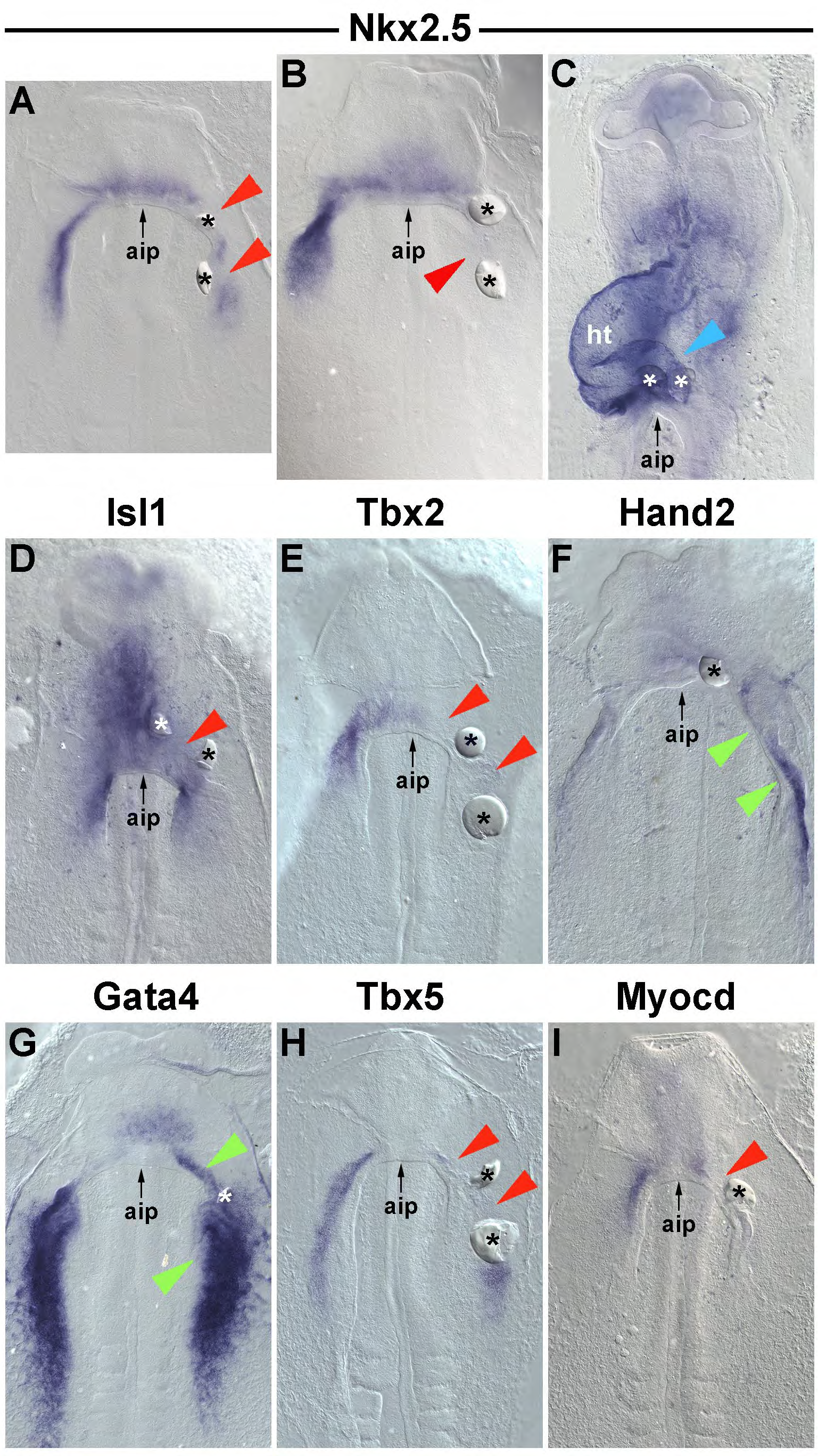
Wnt3a has differential effects on cardiac and lateral mesoderm markers at HH5/6 and HH7/8. Ventral views of embryos with Wnt3a-loaded beads implanted into the lateral/cardiogenic mesoderm at HH5/6 (A), HH7/8 (B, D-I) and HH9/10 (C). After 6 hours of re-incubation, embryos were assayed for the markers indicated on top of each image. Note that *Nkx2.5*, *Isl1*, *Tbx2*, *Tbx5*, *Myocd* were downregulated (A,B,D,E,H,I; red arrowheads); *Hand2* and *Gata4*, two genes also expressed in the trunk lateral mesoderm were upregulated (green arrowheads), consistent with Wnt caudalising the rostral lateral mesoderm. At HH9/10, *Nkx2.5* was refractory (C, blue arrowhead); *Isl1* can still be downregulated (not shown). Abbreviations: aip, anterior intestinal portal; ht, heart.

**Figure 12.**
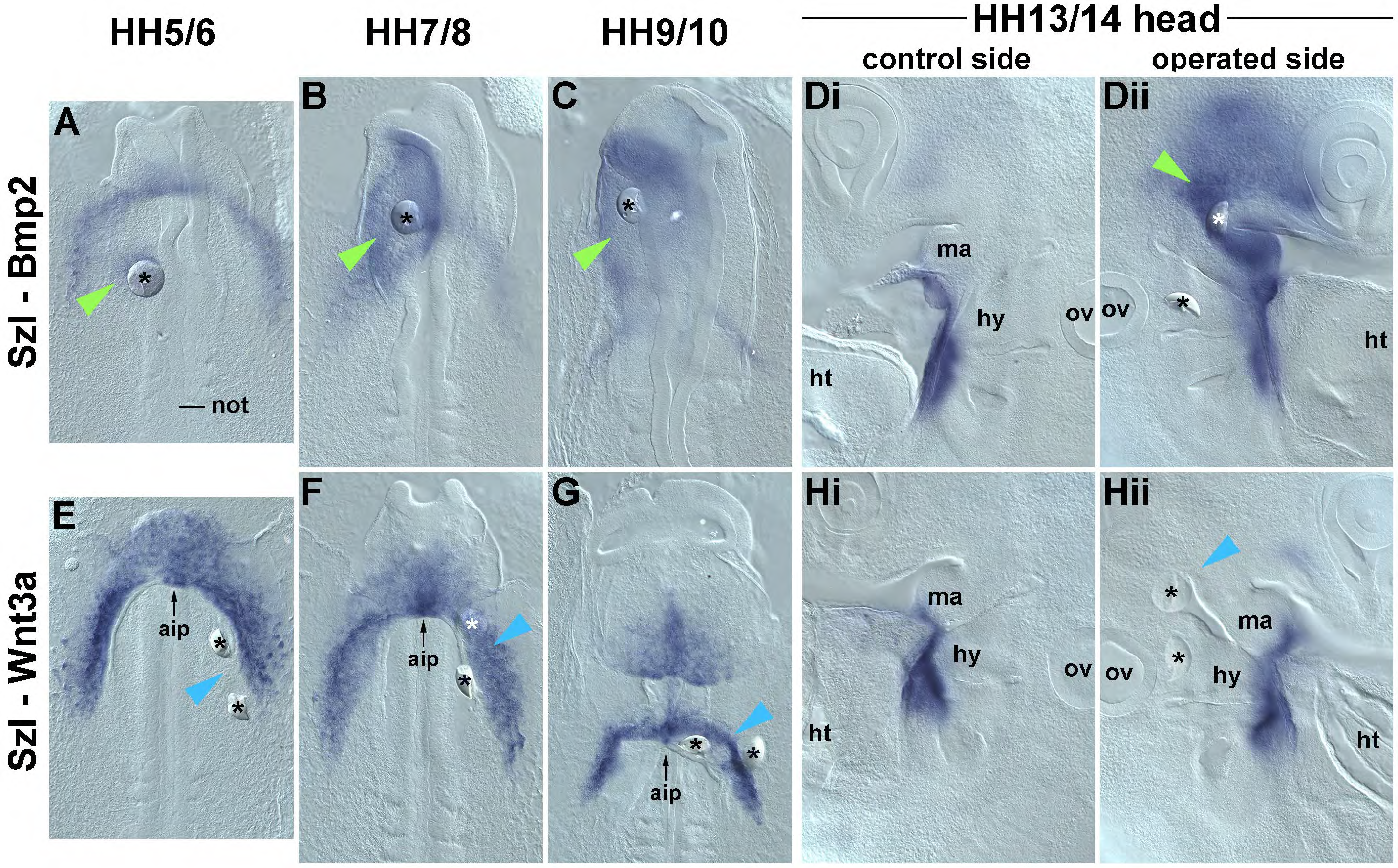
Effect of Bmp and Wnt on the Wnt-inhibitor *Szl*. (A-Dii) Bmp2-loaded beads were implanted into the PHM of HH5/6, HH7/8, HH9/10 and HH13/14 embryos as in Fig.3,4,8-10; embryos were re-incubated for 6 hours. Dorsal views in (A-C), lateral views in (Di,Dii). *Szl* was strongly upregulated at all times. (E-Hii) Wnt3-loaded beads were implanted into the lateral/ cardiogenic mesoderm of HH5/6, HH7/8, HH9/10 embryos as shown in Fig.11, and into the pharyngeal mesoderm of HH13/14 embryos as in (Dii). Ventral views in (E-G), lateral views in (Hi,Hii). Wnt3a did not affect *Szl* expression. Abbreviations as in Fig.3,4,8-11.

**Figure 13.**
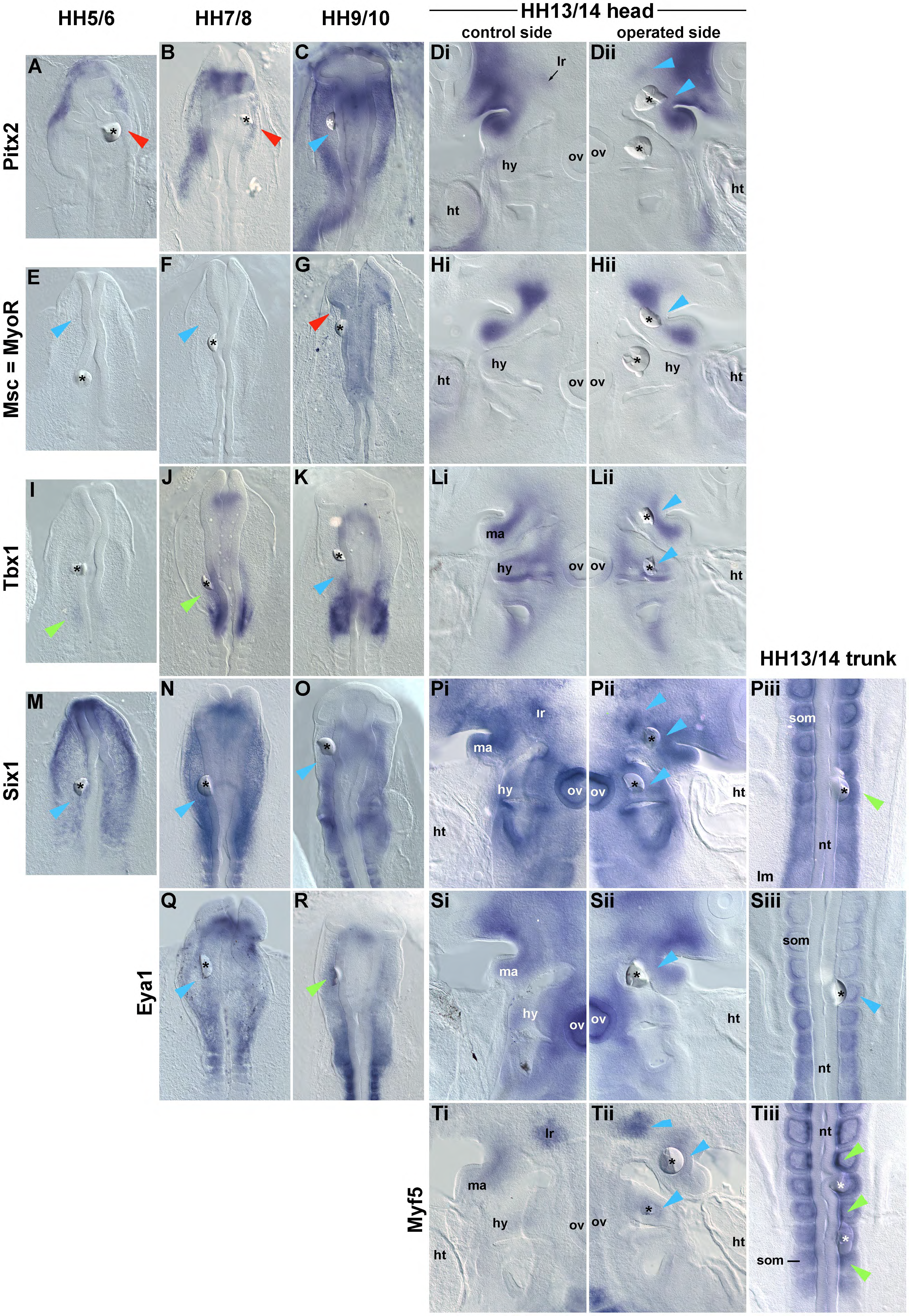
Effect of Wnt3a on markers for the head programme of myogenesis and on generic skeletal muscle markers. Wnt3a-loaded beads were implanted into the PHM of HH5/6, HH7/8, HH9/10 and HH13/14 embryos as in Fig.12E-Hiii. For markers also expressed in the trunk, beads were additionally placed into the somites (Piii,Siii,Tiii). After 6 hours, embryos were assayed for the expression of marker genes shown on the left of the panel. Embryos are displayed in dorsal or lateral views as in Fig.3,4,8-10; annotations and abbreviations also as before. In the head, Wnt3a suppressed the expression of *Pitx2* and *Msc* (A,B,G; red arrowheads), slightly expanded the expression domain of *Tbx1* (I,J; green arrowheads) in a rostral direction, and slightly elevated *Eya1* expression to levels found in the trunk (R, green arrowhead), again consistent with Wnt caudalising the PHM. At stages HH13/14, these markers were unaffected (Dii-Sii; blue arrowheads), as was *Myf5* (Xii; blue arrowhead). In the trunk, Wnt3a slightly elevated the expression of *Six1* and *Myf5* (Tiii, Xiii).

#### 3.5.1. Effect of Wnt on cardiac markers

When Wnt3a-beads were inserted into the cardiogenic region of HH5/6 and HH7/8 embryos and the embryos left to develop for 6 hours, expression of *Nkx2.*5 was downregulated (Fig.11A,B; red arrowheads). However, when implanted at HH9/10, *Nkx2.5* expression was stable (Fig.11C, blue arrowheads), suggesting that cardiac markers are most vulnerable to Wnt at early stages of development. Focusing on stages HH7/8, we analysed the effect of Wnt on *Isl1* (Fig.11D), *Tbx2* (Fig.11E), *Hand2* (Fig.11F), *Gata4* (Fig.11G), *Tbx5* (Fig.11H) and *Myocd* (Fig.11I). We found that Wnt mildly downregulated *Isl1*, and more strongly repressed *Tbx2*, *Tbx5* and *Myocd* (Fig.11D,E,H,I; red arrowheads), corroborating the early Wnt-sensitivity of cardiogenic cells. Yet the 6-hour Wnt exposure led to some upregulation of *Hand2* and *Gata4* (Fig.11F,G, green arrowheads), two of the markers more widely expressed in the lateral mesoderm, including in the trunk. This finding is consistent with a caudalising role of β-Catenin-mediated Wnt signalling.

#### 3.5.2. Effect of Bmp and Wnt on *Szl*

Testing the effect of Bmp and Wnt on the expression of the Wnt inhibitor *Szl*, we found that Bmp very strongly induced *Szl* at all stages examined (Fig.12A-Dii, green arrowheads). In contrast, Wnt did not affect *Szl* expression, neither in the early cardiogenic mesoderm (Fig.12E-G, blue arrowheads) nor in the HH13/14 pharyngeal arches (Fig.12Hii; blue arrowhead). Thus, protection from Wnt signalling is available to cells responding to Bmp at all times. This suggests that the loss of cardiac competence in the PHM is not due to the elevation of Wnt signalling or loss of protection from Wnt.

#### 3.5.3. Effect of Wnt on markers for the head skeletal muscle programme

When Wnt3a-beads were implanted into the PHM, they downregulated *Pitx2* at HH5/6 and HH7/8 (Fig.13A,B; red arrowheads), and prevented the onset of *Msc* at HH9/10 (Fig.13G, red arrowhead). Wnt at times mildly expanded the expression domains of *Tbx1* in a rostral direction (Fig.13I,J; green arrowheads) or upregulated rostral *Eya1* levels to those found more caudally (Fig.13R, green arrowheads), in line with a generic caudalising role of Wnt. However, within the 6-hours timeframe used here, Wnt3a had no effect on *Six1* and *Myf5* expression in the head (Fig.13M-Pii; Fig.13Tii; blue arrowheads) whilst expression in the trunk was slightly upregulated (Fig.13Piii, Tiii; green arrowheads). Taken together, Wnt had a negative effect on the head skeletal muscle programme, largely by suppressing the genes establishing the precursor cells.

### 3.6. The effect of Bmp and Wnt on the cranial signalling landscape

To this end, our experiments suggested that β-Catenin-mediated Wnt signalling suppressed the cardiac as well as the head skeletal muscle programme, in line with its generic caudalising role. However, markers expressed in the caudal PHM (*Tbx1*) or more prominent in the occipital somites (*Lef1, Eya1*) were only mildly expanded in a rostral direction. We therefore turned to *Cyp26C1*, a generic PHM marker as well as a retinoic acid inhibitor; *Bmp2* and *Bmp7*, two genes relevant cardiac development but also signalling from the prechordal plate; the prechordal plate marker *Gsc* the prechordal and notochord marker *Shh*, the forebrain marker *Six3*, and the pharyngeal pouch marker *Fgf8*.

We first, however, tested whether Bmp loaded onto acrylic beads would suppress *Cyp26C1* in the PHM similar to AffiGel blue beads used before (Bothe et al., 2011). Indeed, we found that Bmp suppressed this marker, most strongly at HH5/6 and 7/8 (Fig.14A-C, red arrowheads). The gene was refractory at HH13/14 (Fig.14Dii, blue arrowheads). When Wnt3a-beads were implanted, they similarly suppressed *Cyp26C1* at HH5/6, HH7/8 and HH9/10 (Fig.14E-G, red arrowheads), both in rostral and caudal areas of the PHM. This suppression did not occur at HH13/14 (Fig.14Hii, blue arrowheads). Downregulation was observed already after 2 hours (see Fig.2V-X), identifying *Cyp26C1* as the most Wnt-sensitive marker in our study. When Wnt3a beads were implanted into the lateral/cardiac mesoderm, they slightly downregulated the expression of *Bmp2* and *Bmp7* (Fig.14I,K, red arrowheads). Wnt-beads implanted into the rostral PHM near the prechordal plate however substantially suppressed both Bmps (Fig.14J,L, red arrowheads). Wnt beads also downregulated the prechordal expression of *Gsc* and the expression of *Six3* in the forebrain (Fig.14M,N, red arrowheads), but did not affect *Fgf8* in the pharyngeal endoderm or *Shh* in the prechordal plate and notochord (ST2, not shown). This suggests that Wnt signalling broadly affects the PHM by lifting the protection from caudal retinoic acid signalling, and by affecting the prechordal plate as the Bmp-releasing signalling centre that initiates the expression of *Msc*.

**Figure 14.**
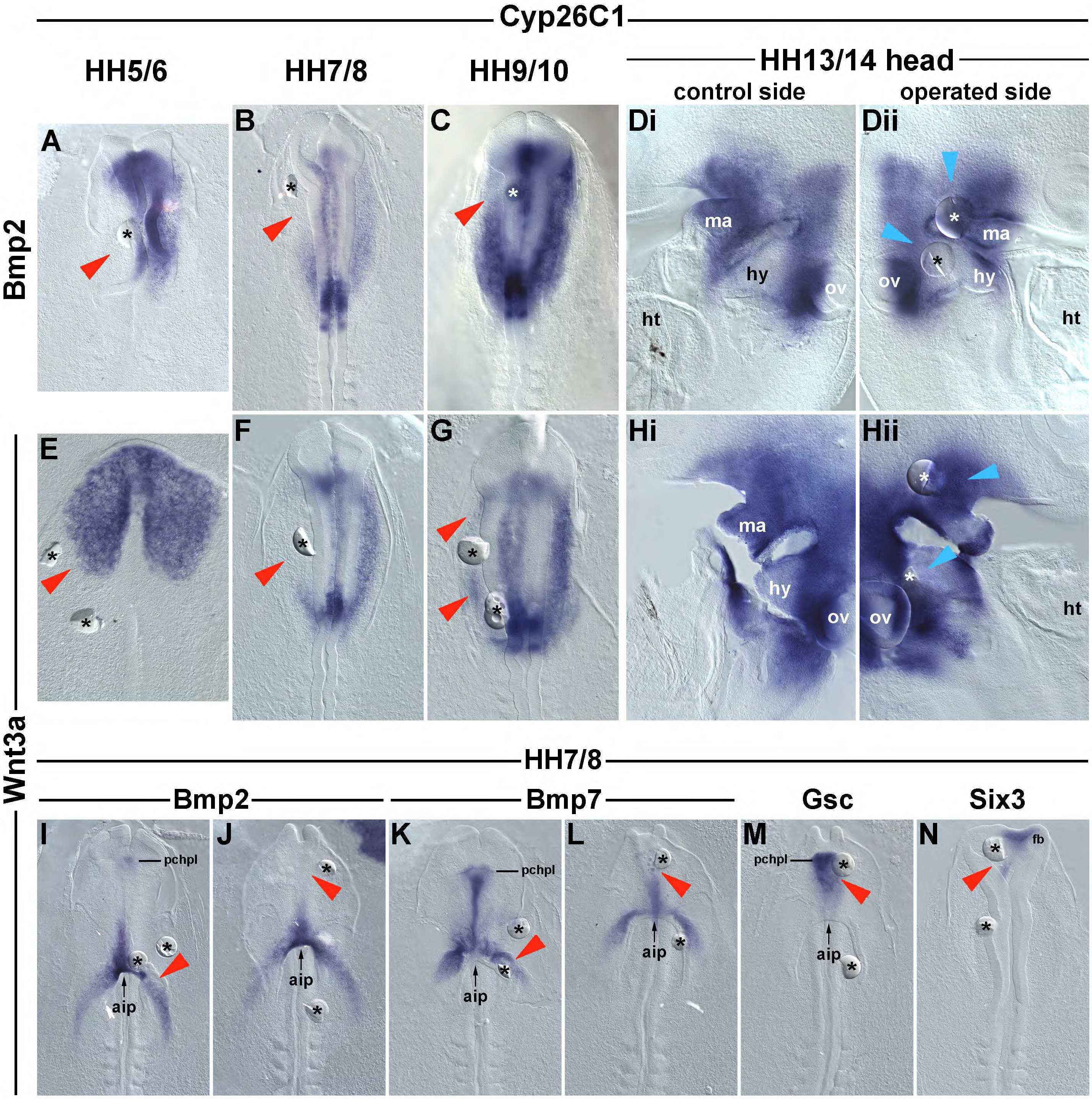
Effect of Bmp and Wnt3a on cranial signalling systems and on the prechordal plate and forebrain. (A-C) Dorsal and (Di,ii) lateral views of embryos treated for 6 hours with Bmp2-loaded beads at the stages indicated on top of the panel, followed by in situ hybridisation for the PHM marker and retinoic acid inhibitor *Cyp26C1*; annotations and abbreviations as before. Consistent with its generic lateralising role, Bmp2 suppressed *Cyp26C1* at stages HH5-10 (A-C, red arrowheads). (E-G) Dorsal and (Hi,ii) lateral views of embryos treated for 6 hours with Wnt3a loaded beads and assayed for the expression of *Cyp26C1*. Between HH5-10, Wnt3a suppressed *Cyp26C1* both in rostral and caudal areas of the PHM (E-G, red arrowheads). (I-M) Ventral and (N) dorsal views of embryos treated for 6 hours with Wnt3a beads at HH7/8; markers are indicated above the images. Abbreviations: aip, anterior intestinal portal; fb, forebrain; pchpl, prechordal plate. When the beads were implanted into the cardiac region (I,K), they slightly downregulated *Bmp2* (I, red arrowhead) and *Bmp7* (L, arrowheads). Whenbeadswereimplanted into the PHM (J,L,M,N), then rostrally located beads downregulated *Bmp2* and *Bmp7* in the prechordal plate (J,L, red arrowheads; compare with I,K). Wnt3a also slightly downregulated *Gsc* in the prechordal plate (M, red arrowhead) and suppressed *Six3* in the forebrain (N, red arrowhead).

**Figure 15.**
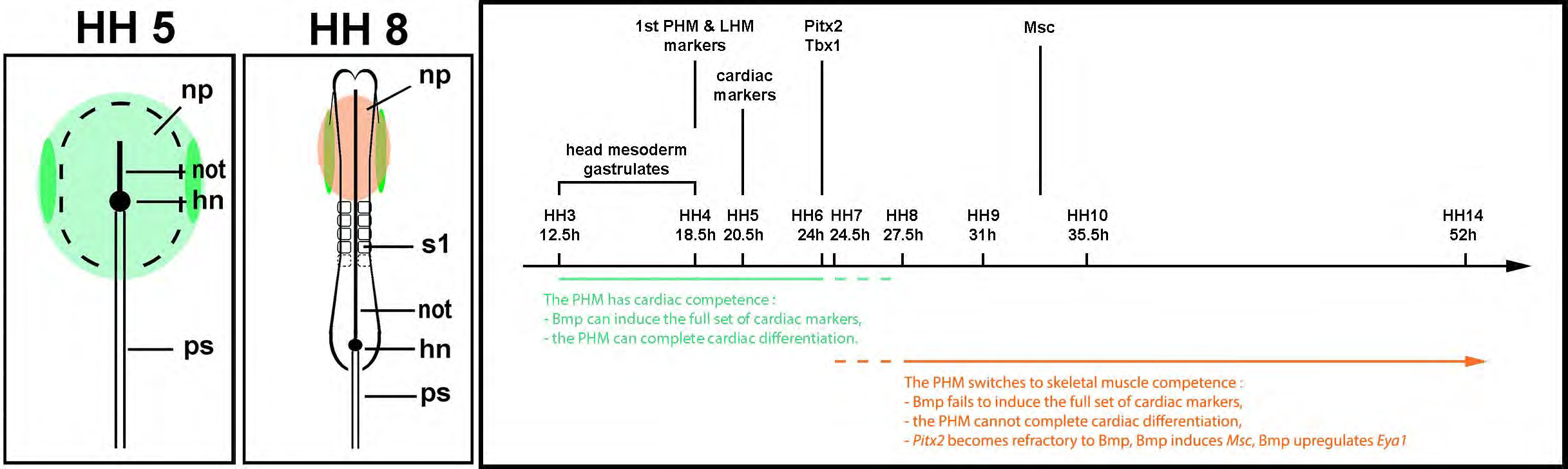
Graphical Summary. The head mesoderm has full cardiac competence until HH5/6, and prolonged smooth muscle competence until HH9/10. In the PHM, the head skeletal muscle programme becomes available as cardiac competence declines when the embryos transit through stages HH7/8. Abbreviations: hn, Hensen’s node; not, notochord; np, neural plate; ps, primitive streak; s1, last-formed somite

## 4. Discussion

Various forms of cell labelling/ cell tracing experiments have mapped the fate of the early gastrulating mesendoderm, i.e. established the normal developmental programmes or trajectories of cells left undisturbed. These studies showed that cells gastrulating from the HH3 chicken primitive streak and settling in a position lateral to the boundary of the overlying neural plate will deliver the primitive heart that starts beating at HH10 ((Camp et al., 2012; Garcia-Martinez and Schoenwolf, 1993); reviewed in (Sendra et al., 2021; Wittig and Münsterberg, 2020)). In contrast, cells that settle in a paraxial position contribute to the skull and to craniofacial skeletal muscles (reviewed in (Schubert et al., 2018). The fate of the cells at the paraxial-lateral interface is somewhat unclear, likely because these cells are multipotent and able to contribute to the SHF/heart, to the heart’s smooth muscle collar, to the great vessels as well as head skeletal muscle.

Exposing the head mesoderm to molecular challenges has led to an enormous body of knowledge regarding the extrinsic factors (signalling molecules, but also tissue tension) and the intrinsic factors (downstream modulators of signals and transcription factors) that control normal development. However, the distinction between cell fate and the often much wider cellular competence or potency, i.e. the ability to engage in a programme, has rarely been addressed. In addition, the timescale of experiments has often led to differing views on the state of cellular commitment or plasticity, with cell specification, i.e. the reversible commitment of cells to a developmental programme, and cell determination, i.e. the irreversible commitment of cells to a developmental programme, not always being distinguished. Here in this study, we aimed to establish whether there is a generic cardiac competence in the head mesoderm, encompassing the paraxial head mesoderm (PHM) that is not fated to contribute to the heart.

### 4.1. At early headfold stages, the PHM is heart-competent

Using a strict 6-hour regime to treat the chicken PHM with the heart-inducing signalling molecule Bmp2, we found that up to early head fold stages at HH5/6, the PHM was able to activate the palette of transcription factors that cooperatively drives cardiogenesis in vitro and in vivo. Moreover, when given time to complete differentiation, the early PHM was able to do so, both when exposed to ectopic Bmp or when grafted into the cardiogenic region of HH5-8 hosts. Thus, the entire head mesoderm is initially heart-competent. Notably, at early head fold stages, the PHM already expresses *Cyp26C1* that protects the tissue from caudalising retinoic acid. Moreover, it just about switched on *Pitx2* and *Tbx1*, two genes later involved in the activation of the skeletal muscle programme ((Bothe and Dietrich, 2006) reviewed in (Schubert et al., 2018)). This suggests that at HH5/6, the PHM is specified but not yet determined to follow its own developmental programme.

### 4.2. The cardiac competence of the PHM fades at early neurula stages

Between HH7-HH10, the ability of Bmp to trigger the cardiac programme in the PHM declined as fewer and fewer of the cardiac genes were able to respond. Moreover, the PHM lost the ability to complete cardiac differentiation. Genes that remained activatable were known Bmp pathway genes and genes associated with general lateral identity both in the trunk and the head mesoderm, indicating that the PHM remained Bmp-sensitive. This suggests that the PHM sheds specifically cardiac competence at early neurula stages. Previous studies had suggested that cardiogenesis can be triggered in the PHM as late as HH13/14 (Tirosh-Finkel et al., 2006). However, these studies treated embryos at HH9/10, just allowing them to develop for a longer time. Moreover, a more limited set of marker genes was analysed, and these markers included general lateral mesoderm markers that can lead to misinterpretation. Yet these data and ours can be reconciled: we found that at HH9/10, Bmp can still trigger the expression of *Isl1*, *Nkx2.5* and *Tbx2*, genes that, rather than being heart-specific, are expressed in various types of precursor cells, both in the endoderm and the ectoderm. One of the mesodermal cell types is smooth muscle, and we found that smooth muscle actin (*Acta2*) as well as the smooth muscle regulator *Csrp2* can be activated by Bmp as late as HH10. This suggests that smooth muscle competence is retained longer than cardiac competence, in line with the requirement of the head mesoderm to finalise the cardiac outflow tract by adding its smooth muscle collar (Wang et al., 2017). At HH13/14, whilst Bmp cannot trigger expression of cardiac genes in the PHM within the 6-hours timeframe, it can dorsally expand the expression domain of *Isl1*. This phenotype was also reported for the long-exposure experiments (Tirosh-Finkel et al., 2006). *Isl1* remains strongly expressed in multipotent cells in the SHF. Moreover, Bmp plays a role in recruiting cells from the SHF into the heart and into differentiation. Taken together, this suggests that at this late stage ectopic Bmp may dorsolaterally expand the area in the subpharyngeal mesoderm from which SHF cells are being recruited, but it will not induce cardiogenesis in the PHM.

### 4.3. At early neurula stages, the developmental competence of the PHM shifts to the head skeletal muscle programme

As embryos progressed from HH5/6 to HH7/8 and HH9/10, the response of the PHM to Bmp changed markedly. Initially, Bmp suppressed a wide palette of PHM marker genes including *Cyp26C1*, genes associated with the head programme of skeletal muscle formation (*Pitx2* and *Tbx1*), and genes generally involved in myogenesis (*Six1* and *Eya1*; reviewed in (Schubert et al., 2018)). These genes became more and more resistant to Bmp, with *Eya1* being upregulated by Bmp at HH9/10. Moreover, from HH7/8 onwards, Bmp activated *Msc*, a further upstream regulator of head skeletal myogenesis. This suggests that at the time cardiac competence fades, skeletal muscle competence is established. This however did not lead to a premature entry into differentiation. Skeletal muscle differentiation in the head does not commence before HH13/14 (Meireles Nogueira et al., 2015), and both for head and trunk skeletal muscle differentiation, Bmp is a strong inhibitor ((Tzahor et al., 2003; von Scheven et al., 2006a); reviewed in (Schubert et al., 2018)). Moreover, Msc, whilst upregulating *Myf5* and *Myod* at later stages (Moncaut et al., 2012), mainly acts as a transcriptional repressor (Harris et al., 2015), suggesting a scenario whereby the head skeletal muscle precursor genes establish myogenic competence but keep differentiation at bay to allow the expansion of the precursor pool first.

### 4.4. The triple role of Wnt in suppressing head mesodermal programmes

Previous studies suggested that from HH7/8 onwards, the cardiac programme may be repressed in the PHM owing to rising levels of β-Catenin-mediated Wnt signalling (Tzahor and Lassar, 2001). We thus expected that ectopic Wnt3a application would promote the expression of PHM genes. However, this was not the case: Wnt3a suppressed the expression of most PHM and skeletal muscle associated genes in the same fashion as it repressed cardiac genes. At the same time, expression of genes representing more caudal positional values were upregulated and rostrally expanded. This suggests that Wnt facilitates the caudalisation/posteriorisation of the mesoderm, both in the paraxial and the lateral region ((Bressan et al., 2013; Marvin et al., 2001); this study). Compared to the strong lateralising effects of Bmp, the Wnt-induced rostral expansion of caudal markers was more subtle, probably because Wnt inhibitors are expressed in Hensen’s node (*Crescent, Sfrp1, Dkk1*), the early endoderm (*Crescent, Sfrp1*), the neural plate/ neural tube next to the PHM (*Sfrp1, Sfrp2, Sfrp3, Dkk2*), the notochord (*Dkk1*) and the lateral and cardiogenic mesoderm (*Szl*; see http://geisha.arizona.edu/geisha/), all protecting rostral tissues from the caudalising influence of Wnts produced in the primitive streak.

In addition to its generic caudalising effect, at all the early stages from HH5-10 Wnt strongly suppressed *Cyp26C1*, both in rostral and in caudal territories of the PHM. Cyp26C1 protein is a retinoic acid inhibitor and the first factor to distinguish the PHM (Bothe and Dietrich, 2006). Moreover, whilst the Wnt-expressing primitive streak moves caudally, the expression domain of the key enzyme to produce retinoic acid, Aldh1a2 (Raldh2) expands rostrally to specify the cardiac atria and to separate atria and ventricle (Hochgreb et al., 2003), reviewed by (Duong and Waxman, 2021), suggesting that the control of retinoic acid signalling may be a more critical step in HH5-10 head mesoderm development. Indeed, retinoic acid has been shown to suppress the head skeletal muscle precursor markers in the PHM, with a more prominent effect on the rostral markers *Pitx2* and *Msc* than on *Tbx1* (Bothe et al., 2011). Thus, much of the caudalising effect of Wnt on the PHM may be indirect and via retinoic acid, and may require more than 6 hours to unfold.

Notably, we found that Wnt signalling, whilst affecting Bmp expression in the lateral/ cardiac region only marginally, substantially suppressed Bmp signalling molecules in the prechordal plate. The prechordal plate marker *Gsc* and the forebrain marker *Six3* were also downregulated, in line with work conducted in the zebrafish that established prechordal plate and forebrain genes as immediate Wnt targets (Green et al., 2020). Bmp released by the prechordal plate is required for the induction of *Msc* (Bothe et al., 2011). Thus, while we cannot rule out a direct suppression of *Msc* by Wnt, Wnt may prevent *Msc* activation via suppression of prechordal plate signalling.

Previous studies established that Wnt signalling suppresses skeletal muscle formation in the head mesoderm whilst promoting myogenesis in the trunk, with Wnt acting directly in the activation of Mrf family members in the somite (reviewed in (Schubert et al., 2018)). *Mrf* gene expression in the head does not commence before HH13/14 (Meireles Nogueira et al., 2015). Yet within the time frame of 6 hours, cranial *Myf5* or *Myod* expression did not change in response to Wnt at these stages (this study). In past studies, ectopic Wnt was applied at the early stages when we found that Wnt would suppresses head skeletal muscle precursors genes, and the embryos were allowed to develop for a long time (Tzahor et al., 2003). It thus seems likely that in all studies, Wnt prevented cranial myogenesis by interfering at the level of the upstream regulators.

### 4.5. The shift from cardiac to skeletal muscle competence is intrinsic to the PHM

Our RNAseq screen revealed *Szl* as one of the most Bmp-dependent genes at HH7/8. Szl is a Crescent-and Sfrp1/2/5-related Wnt inhibitor that is specifically expressed in the early cardiogenic mesoderm, offering protection from heart-suppressing Wnt. We found that Bmp strongly and quickly upregulated *Szl* in the PHM at all stages investigated. Thus, Wnt protection was always induced, and hence the loss of cardiac competence in the PHM was not due to the overriding effects of Wnt. Likewise, Bmp drove the expression of lateral ectoderm and endoderm markers towards the midline at stages when the PHM had lost the ability to complete cardiogenesis, suggesting that the signalling landscape for cardiogenesis can still be established. Together, this suggests a Wnt-independent loss of cardiac competence in the PHM and, whilst not excluding other extrinsic factors, points to an intrinsic control mechanism. This will require further investigation, but several lines of evidence support this idea: Msc is expressed during formation of the atrioventricular conduction system (von Scheven et al., 2006b), where it physically interacts with Gata4 to repress *Cx30.2* (Harris et al., 2015). Msc may well interact with Gata4 or other proteins earlier to repress the cardiac programme in the PHM. Moreover, as Msc binds to E-boxes similar to those recognised by MyoD (MacQuarrie et al., 2013), and Msc and MyoD binding sites have been predicted for the promoters of several cardiac genes (P. Mendes Vieira and S. Dietrich, unpublished observations), it may repress target genes directly. In a similar vein, Six1 contributes to the upregulation of pharyngeal Fgf8 expression which in turn upregulates *Tbx1* and *Msc* in the mesoderm (Bothe et al., 2011; Guo et al., 2011). Yet it may well have additional roles when it begins to be expressed in the early PHM.

### 4.5. Outlook

During vertebrate evolution, aspects of the skeletal muscle programme were recruited into the head mesoderm twice: first the effector genes required for muscle striation were brought under the control of smooth/cardiac upstream regulators, thereby creating powerful, yet tireless cardiac muscle ((Brunet et al., 2016), reviewed by (Poelmann and Gittenberger-de Groot, 2019; Simões-Costa et al., 2005). Thereafter the core skeletal muscle regulators were recruited, diverting a subset of the head mesoderm to a skeletal muscle programme that allows voluntary control over the cranial openings (reviewed in (Schubert et al., 2018)). As PHM cells are relinquishing cardiac competence, they are potentially in a fragile state as the possibility to engage in skeletal muscle formation becomes available, but genes that enforce myogenesis are not expressed for a prolonged period of time. Pathological heart failure also creates a fragile state in which skeletal muscle-associated genes are de-repressed, most prominently Six1 which upregulates miR-25 that downregulatesSerca2a(Ohetal.,2019).This suggests that there is a cardiac-skeletal muscle antagonism that requires tight control. Unravelling this mechanism will shed light onto normal development and will give deeper insight into cardiac disease.

## Supporting information

Highlights

## Acknowledgements

We thank Soraya Idris-Anderson and Ashley Haywood for their assistance to the project, and the IBBS section Epigenetics and Development for insightful comments. The work was funded by an UoP grant, a UoP funded PhD studentship to M. Wolton, an Erasmus Scholarship to N. Murciano and a BSDB Gurdon studentship to P. Mendes Vieira.

