## Supplementary material for "Cardiac competence of the paraxial head mesoderm fades concomitant with a shift towards the head skeletal muscle programme": Highlights

- Stage-dependent responses to Bmp indicate that initially the paraxial head mesoderm is bestowed with full cardiac competence. This cardiac competence persists until early head fold stages. Thereafter, cardiac competence is replaced by skeletal muscle competence.
- Wnt signalling caudalises (posteriorizes) the head mesoderm, suppressing both the cardiac and the head skeletal muscle programme. Wnt signalling intercepts these programmes by suppressing the key upstream regulators of both programmes. In the PHM, Wnt in addition suppresses signalling from the prechordal plate and removes the protection from retinoic acid signalling.
- Our data are consistent with the idea that cardiac and smooth muscle competence is the evolutionary and developmental ground state of the vertebrate head mesoderm.
