## Supplementary material for "Cardiac competence of the paraxial head mesoderm fades concomitant with a shift towards the head skeletal muscle programme": Legends Suppl material

### Legends for Supplementary Figures and Tables.

#### Supplementary Figure SF1. Schematic representation of previous studies that exposed the chicken PHM to Bmp and Wnt in comparison with our approach.

Top of panel: Average developmental times and embryo morphology (dorsolateral views) are plotted onto a time line. Standard bead implantation sites to target the paraxial head mesoderm (PHM) in our study are also shown (turquoise circles), deviations from these sites are explained in the main text. Abbreviations: epi, epiblast; hn, Hensen’s node; ht, heart; not, notochord; np, neural plate; ps, primitive streak; r1/2, rhombomeres 1-2; s1, 1^st^ somite; tb, tail bud.

Beneath in black: period of head mesoderm gastrulation and onset of expression for key markers in the cardiogenic mesoderm (CM) and paraxial head mesoderm (PHM).

Pink: time period/ developmental stage covered and results obtained from previous short-term Bmp exposures. Red: developmental stage covered and results obtained from experiments in which the PHM was exposed to Bmp continuously and for a long time. Blue: developmental stage covered and results obtained from experiments in which the cardiogenic mesoderm was exposed to Wnt continuously and for a medium time-range. Green: approaches and results from this study; ON, overnight exposure.

Note that manipulations performed at HH3-4 in the first instance affect the migration of head mesodermal cells. The outcome of manipulations at later stages depends on the experimental design. Short-term exposure allows to study immediate effects. Long-term, continuous exposure prevents the system from rightening itself. It allows to study downstream effects, but does not allow to determine when this effect was created, and how.

#### Supplementary Figure SF2. Within the 6-hours timeframe, Bmp does not activate genes controlling myocardial differentiation.

Beads loaded with 6.41µM Bmp2 were implanted into the PHM of HH5/6, HH7/8 and HH9/10 embryos as shown in Fig.4. After 6 hours, the embryos were analysed for the expression of genes controlling myocardial differentiation as shown on the left of the panel. Moreover, Bmp2-beads were used on HH5/6 embryos at 19.23µM to test whether a higher concentration can force differentiation within 6 hours. However, this was not the case (blue arrowheads), the only response was an upregulation of *Mef2c* in emigrating neural crest cells (C, green arrow).

#### Supplementary Table ST1. Probes used for in situ hybridisation, n-numbers and phenotypes for bead grafting experiments.

##### A. In situ probes and responses of marker genes to 6-hours of treatment with Bmp- and Wnt-loaded heparin-coated acrylic ‘white’ beads.

Columns A-E contain the description of each marker with key expression domains and hyperlinks to published probes and expression patterns. Columns H-L, O-S, show the n-numbers per Bmp and Wnt treatment and developmental stage at the time of bead implantation. Bmp beads were all implanted into the paraxial head mesoderm (PHM) or the flank somites. At HH5/6, 7/8, 9/10 (Columns O-Q), Wnt beads were implanted into the PHM, the rostral PHM or the cardiogenic mesoderm (CM) as indicated in the individual cells. Columns G,N indicate the figures showing a representative specimen.

Unchanged expression is denoted as ‘wt’ wildtype, and cells are labelled blue. For upregulated expression, cells are marked green, for downregulation, cells are marked red.

Abbreviations: CM, cardiogenic mesoderm; pchpl, prechordal plate; PHM, paraxial head mesoderm.

Total number of specimen on this sheet: 705.

##### B. Negative controls, comparison of different Bmp2 concentrations and different bead types; 6-hour and overnight exposure.

Columns A,B indicate the marker and stages that were investigated; Column C refers to the respective figure. Columns E-I show the n-numbers for the controls and the Bmp2-loaded white beads (6 hours exposure), Columns K-O show the corresponding data for Affi-Gel blue agarose beads, Columns Q-T show the n-numbers for overnight experiments. Colour code of cells as in (A).

Total number of specimen not counted before: 145.

##### C. Comparison of different Bmp ligands.

Columns A,B indicate the marker and stages that were investigated; Column C refers to the respective figure. Columns E-I show the n-numbers for the controls and the different Bmps loaded onto white beads. Colour code of cells as in (A).

Total number of specimen not counted before: 51.

##### D. Time course to monitor the onset of responses to Bmp and Wnt.

Columns A-D indicate the marker gene; the ligand, its concentration and the bead-type used; the developmental stages investigated and the Figure showing representative specimen. Columns F,G,H display the n-numbers for 2 hour, 4 hour and 6 hours exposure, respectively. Colour code of cells as in (A).

Total number of specimen not counted before: 38. Total number of beaded specimen in this study: 939.

#### Supplementary Table ST2. N-numbers for the tissue grafting experiments.

Displayed are the number of embryos with myocardial differentiation in the graft as revealed by the MF20 antibody, compared to the total number of embryos that had the graft at the site indicated. The percentage of embryos with myocardial differentiation has also been calculated and a corresponding colour scheme has been applied. For all graftings, the intended target site was the cardiogenic mesoderm (CM); the hit/retention rate for HH5/6 hosts was 30:54 = 56%, and for HH7/8 hosts, it was 42:46 = 91%. A total of 100 graftings were analysed.

Abbreviations: CM, cardiogenic mesoderm; oft, outflow tract; PHM, paraxial head mesoderm.

#### Supplementary Table ST3. Genes identified in the HH7/8 RNAseq screen as meeting or exceeding the threshold for Bmp-dependency.

Genes that not only showed reduced expression upon 6 hours of Bmp suppression (=loss-of-function approach used for the RNASeq screen) but were induced or upregulated by 6 hours of Bmp on beads (=gain of function experiment) are shown in strong green. Genes that were only upregulated earlier at HH5/6 are shown in light green. Genes for which Bmp was not sufficient to induce expression are shown in blue. These genes are likely indirect Bmp targets.
