## Supplementary figures and images for "Cardiac competence of the paraxial head mesoderm fades concomitant with a shift towards the head skeletal muscle programme"

### Suppl Fig SF1

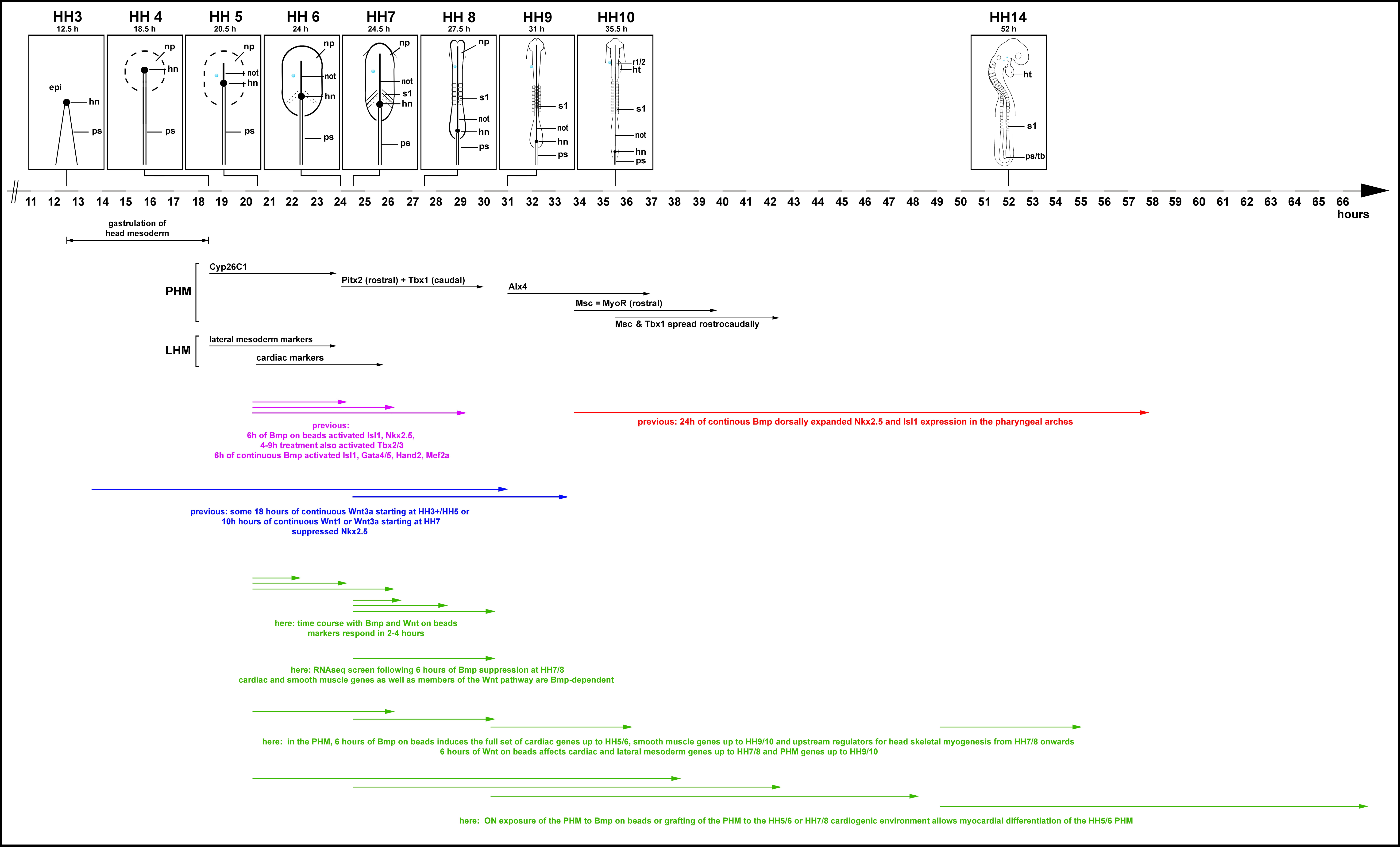

### Suppl Fig SF2

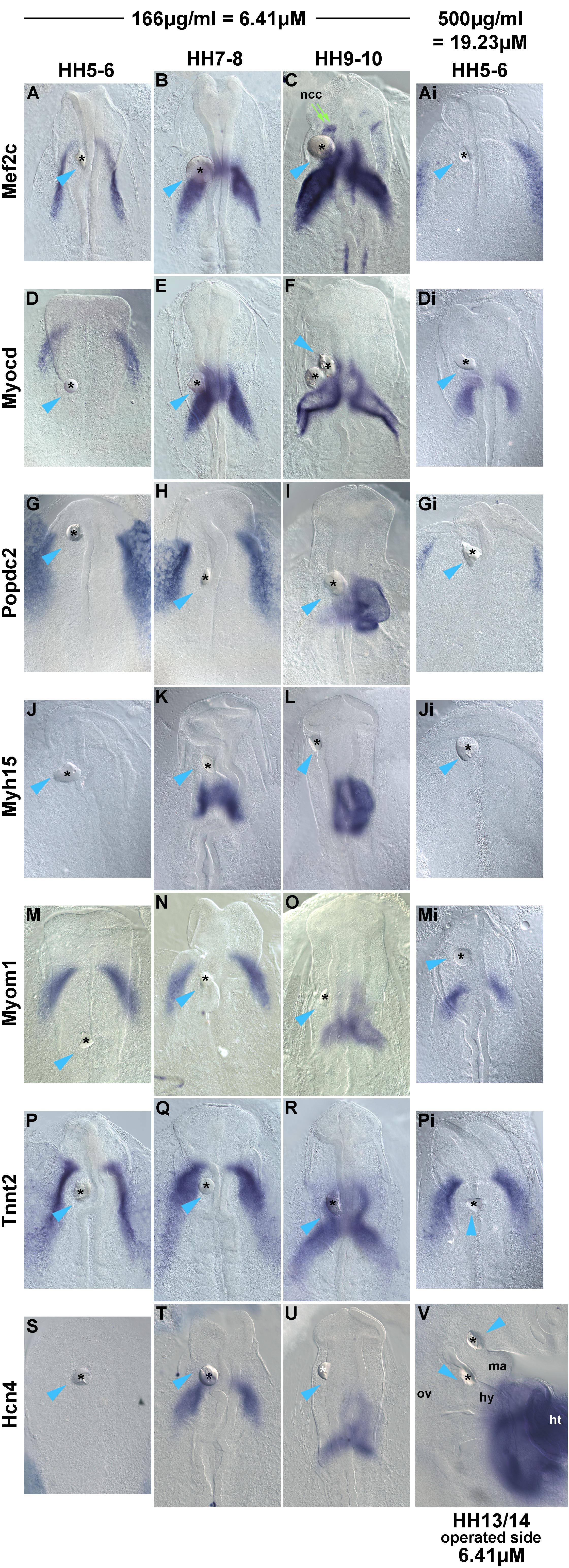
